# Development and Validation of a Continuous Real-Time Optical Sensor for Indocyanine Green Clearance Measurement During Ex-Vivo Perfusion of Human Livers

**DOI:** 10.1101/2025.11.05.686646

**Authors:** Edmund N.J. Derwent, Charles W.G. Risbey, Anita Niu, Paul Yousif, Nidula Fonseka, Spencer Curry, Chloe Seow, Isaac Ng, Geoffrey W. McCaughan, Michael Crawford, Carlo Pulitano, Daniel Babekuhl

## Abstract

Liver transplantation remains the only curative treatment for end-stage liver failure, yet its impact is constrained by organ shortages and graft non-utilisation. Machine perfusion (MP) enables ex-vivo assessment of donated livers; however, existing viability criteria rely on intermittent sampling, reducing temporal resolution and accuracy. Indocyanine green (ICG), a clinically validated dye cleared exclusively by hepatocytes, provides a continuous index of hepatic function beyond initial injury. Accordingly, we present a non-invasive, clamp-on optical sensor that enables continuous, real-time quantification of ICG clearance during MP. The sensor consists of a clamp-on module with an 808nm laser and phototransistor connected to a microcontroller-based unit and computer for real-time plotting. The raw phototransistor signal was linearised to a unitless absorbance signal proportional to perfusate ICG; bi-exponential fitting yielded plasma disappearance rate (PDR_bi_, %/min) and the 15-minute residual fraction (R_15_). Across 10 whole and 3 split human livers (45 boluses; 13 paired with spectrophotometry), the sensor closely matched spectrophotometric measurements (mean R^2^ = 0.994; range 0.983-0.999). The sensor resolved expected physiological trends: ICG clearance increased with temperature (PDR_bi_: 8.2%/min (subnormothermic MP) to 22.6%/min (normothermic MP) (n=4); 9.3%/min (32°C) to 11.9%/min (36°C) (n=1)). The sensor’s continuous signal traces also revealed early mixing dynamics and medication-related effects that are missed by intermittent sampling. This optical sensor enables accurate, real-time monitoring of ICG clearance during ex-vivo perfusion. The ex-vivo setting is uniquely positioned to validate ICG clearance models and enhance clinical interpretation.

## Introduction

Liver transplantation remains the only curative treatment for end-stage liver disease, but its success is limited by the scarcity of viable donor organs (Devarbhavi et al., 2023; Girgenti et al., 2020). Machine perfusion (MP), in which oxygenated perfusate is circulated through an organ ex-vivo, has emerged as a promising strategy to extend the preservation time of livers to up to 24 hours and directly assess graft function before transplantation. Despite these advances, objective and reliable criteria for determining graft viability during perfusion remain elusive. Traditional biochemical markers such as lactate and bilirubin clearance are still widely used, but their correlation with post-transplant outcomes is inconsistent, and no objective method for real-time assessment currently exists (Brüggenwirth et al., 2020; Schlegel et al., 2019). Indocyanine green (ICG), a dye exclusively cleared by hepatocytes, provides a dynamic and mechanistic assessment of liver function and has been used clinically for decades to predict hepatic failure risk. More recently, its feasibility for assessing graft function during ex-vivo perfusion has been demonstrated; however, these approaches rely on intermittent manual sampling and offline spectrophotometric analysis, limiting temporal resolution and clinical utility (Lau et al., 2024; Schurink et al., 2025).

ICG is a water-soluble tricarbocyanine dye that binds almost exclusively to plasma proteins upon intravenous administration and is taken up specifically by hepatocytes (Levesque et al., 2016). It is excreted into bile without undergoing enterohepatic recirculation, with approximately 97% of the administered dose recovered in bile (Noitumyae, 2021; Wheeler et al., 1958). Clearance requires uptake by OATP1B1 and OATP1B3 transporters, cytosolic binding to proteins such as glutathione S-transferases, and excretion via MRP2 into the bile canaliculi (Alam et al., 2018; Fan et al., 2020; Kalliokoski and Niemi, 2009; Köller et al., 2021). These transporters are expressed along the basolateral membranes of hepatocytes in all three hepatic zones – periportal (zone 1), midzonal (zone 2), and centrilobular (zone 3) (Kalliokoski and Niemi, 2009). As a result, ICG clearance reflects integrated hepatocellular metabolic and excretory function across the entire liver, offering a key advantage over metabolic markers such as lactate, which is predominantly cleared by zone 1 hepatocytes and can remain normal despite downstream necrosis (Ferrigno, 2015).

ICG exhibits strong absorbance in the near-infrared range (peak λ ≈ 805 nm) and emits fluorescence at ∼835 nm, making it well suited to optical detection methods (Destro and Puliafito, 1989; Vos et al., 2014). These spectral characteristics, combined with its excellent safety profile (Gasperi et al., 2016), rapid clearance, and hepatic specificity, have established ICG as a clinically validated tool for pre- and intra-operative liver assessment and vascular imaging (Fan et al., 2020). Importantly, its optical properties enable both absorbance and fluorescence based sensing modalities, allowing dynamic, non-invasive monitoring of liver function.

Despite its advantages, current methods for ICG clearance analysis during machine perfusion rely on intermittent blood sampling and offline spectrophotometric analysis, which limit temporal resolution and require sustained operator attention (Lau et al., 2024). The short plasma half-life of ICG, typically less than 5 minutes, makes clearance rates highly sensitive to small sampling errors. Lau *et al*. demonstrated that ICG plasma disappearance rate (PDR) and retention rate (R_15_) correlate with graft survival during LT-NMP, but identified the lack of continuous monitoring as a key limitation, with samples collected at 2-30 minute intervals (Lau et al., 2024). As such, this study aimed to address these limitations through the design, development, and validation of a continuous, optical sensor for real-time tracking of ICG kinetics to enable uninterrupted graft assessment during ex-vivo perfusion.

## Methods

### Study Design and Endpoints

This study is a prospective, single-arm feasibility study with the objective to develop and validate a non-invasive, in-line, real-time optical sensor to measure ICG clearance during ex-vivo perfusion of human liver grafts (Figure 1). Human livers declined for transplantation but consented for research were utilised. Ethical approval was attained prior to commencement from the Sydney Local Health District (SLHD) Human Research Ethics Committee (HREC) (approval number X18-0523 and HREC/18/RPPAH/748).

**Figure 1.**
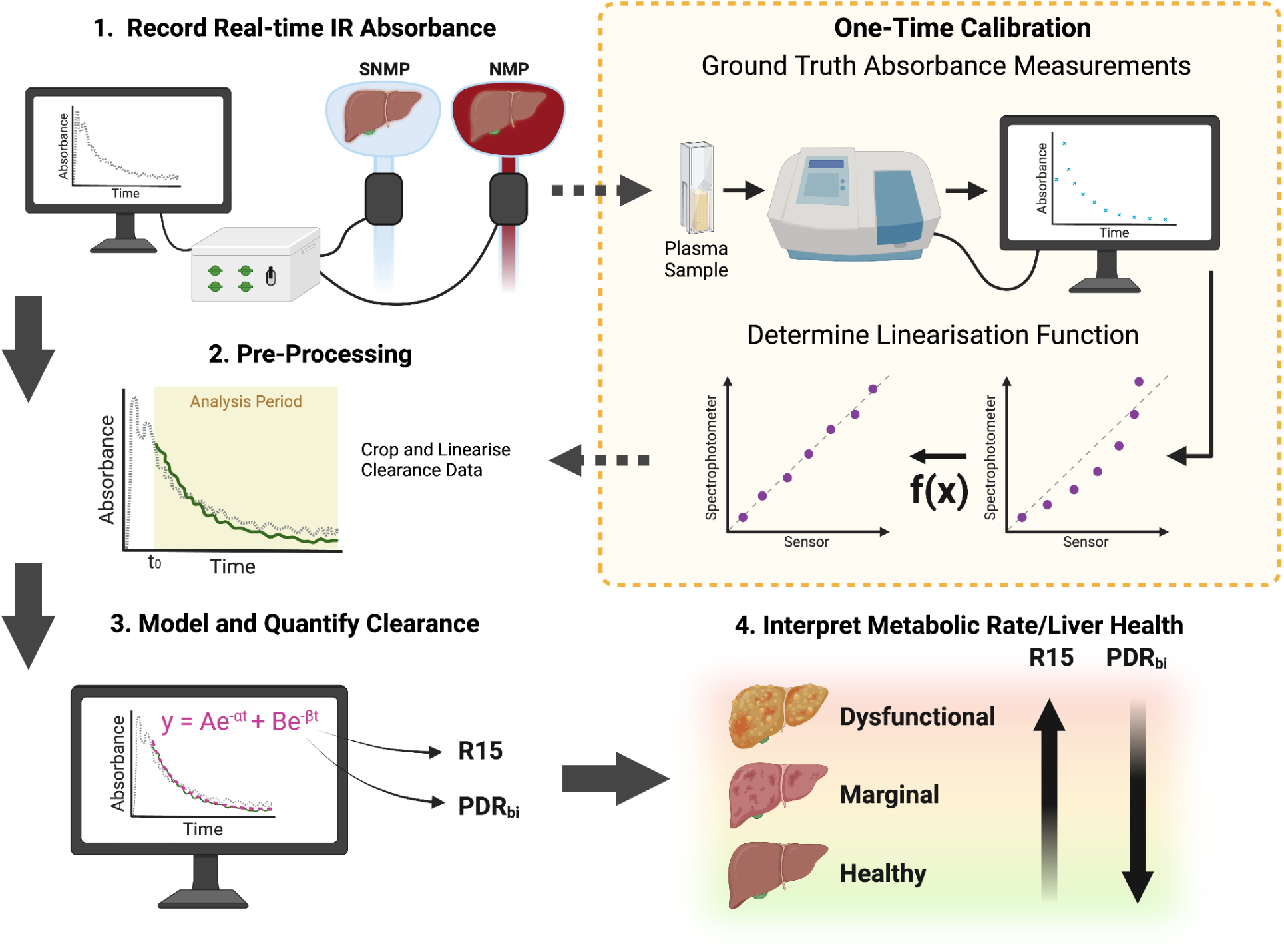
Summary of the ICG sensor approach. A clamp-on sensor records real-time ICG (808nm) absorbance during liver perfusion (SNMP or NMP). A one-time calibration aligns sensor output with ground-truth spectrophotometer measurements. Clearance curves are then fitted to extract key metrics (R15, PDR_bi_), which are used to interpret liver metabolic function and health. Created in BioRender. Derwent, E. (2025) https://BioRender.com/6to7cqt

Normothermic machine perfusion (NMP) was performed using the Extended Organ Perfusion System (EOPS, Aeternum Life Technologies) (Babekuhl et al., 2025). The setup incorporated dual inflow perfusion via the hepatic artery (HA) and portal vein (PV) with continuous blood gas automation, temperature control, and metabolic support using a human red blood cell and plasma based perfusate. The ICG sensor was clamped onto perfusion tubing carrying fully oxygenated blood (oxygen saturation > 98%).

Sub-normothermic machine perfusion (SNMP) was conducted using a modified version of the COARO Hypothermic Oxygenated Machine PErfusion (HOPE) system with dual HA and PV perfusion (Risbey et al., 2025, 2024). Perfusate composition was based on Williams Medium E supplemented with 20% albumin concentrate. The ICG sensor was clamped onto perfusion tubing carrying fully oxygenated perfusate. Livers were transferred between independently controlled SNMP and NMP platforms to enable direct comparison of metabolic clearance rates across thermal states and perfusate compositions.

The primary endpoint was validation of the sensor against the established ICG clearance protocol described by Lau *et al*. via comparison with ground truth spectrophotometry (R^2^) (Lau et al., 2024). Secondary endpoints included extraction of ICG clearance kinetics (plasma disappearance rate (PDR_bi_), and 15-minute retention (R_15_)) and evaluation of the sensor’s ability to detect temperature-dependent differences in clearance between SNMP and NMP perfusion as well as between 32 °C and 36 °C. The overall workflow of the sensor measurement, calibration, and analysis is summarised in Figure 1.

### Sensor Design

#### Physical Design and Layout

The system comprises three modular components: a host computer for data analysis and visualisation, a control unit for data acquisition (housing the microcontroller, laser diode driver and phototransistor sensitivity adjustment), and a clamp-on module designed to interface with transparent medical-grade tubing (Tygon ND-100-65 tubing - ¼″ Internal Diameter, ⅜″ Outer Diameter; Saint-Gobain, France) (Figure S1A). Two iterations of the sensor were developed and tested (versions v1.0 and v1.1). Both employed the same optical configuration, differing slightly in electronic architecture: v1.0 used voltage-driven laser diodes, whereas v1.1 implemented constant-current driven laser diodes to improve optical stability. Inside the module, a 500mW 808nm laser diode (Hangzhou, China) is aligned directly opposite a DFROBOT FIT0180 (Shanghai, China) phototransistor, forming a simple absorbance transmission-mode optical path (Figure 2A, Figure S1B).

**Figure 2:**
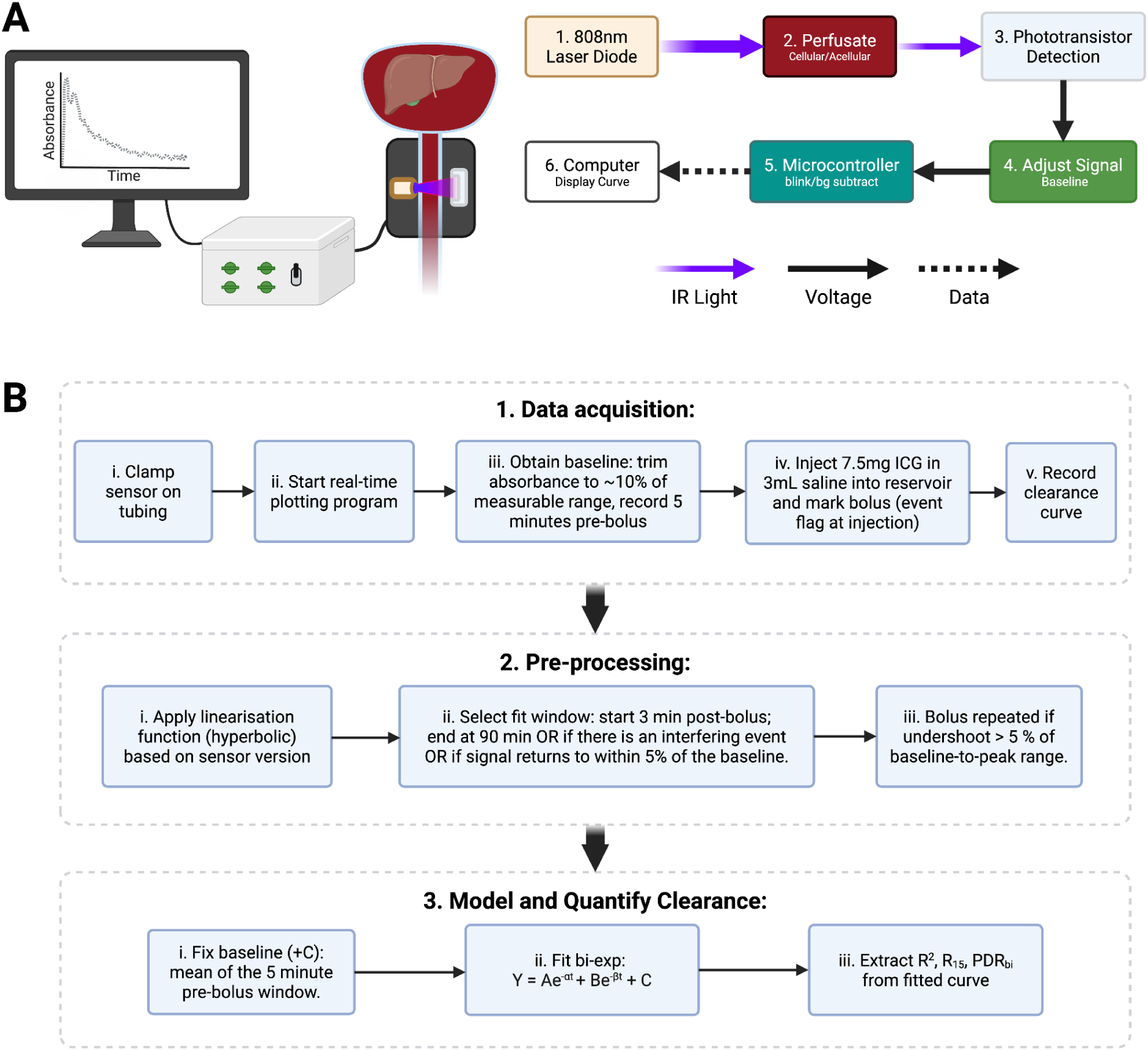
**(A) Clamp-on ICG sensor overview and operating principle.** An 808 nm laser and opposing photodiode measure light attenuation through perfusate according to the Beer-Lambert law, providing a continuous absorbance signal proportional to ICG concentration. The signal is baseline-adjusted and background-corrected before being streamed for real-time visualisation, and saved for curve analysis. **(B) Overview of Signal Processing Pathway.** Schematic representation of the data handling workflow for ICG clearance analysis. Raw sensor absorbance data is collected during perfusion (Data Acquisition), linearised according to the sensor version (Pre-processing), and a bi-exponential decay model is fitted (Curve Fitting). The process yields quantitative clearance metrics: 15-minute retention ratio (R_15_), and plasma disappearance rate (PDR_bi_). Created in BioRender. Derwent, E. (2025) https://BioRender.com/mqsamlh

The control unit drives the laser using an LM317-based constant-current circuit, with coarse (500 kΩ) and fine (10 kΩ) linear potentiometers for manual baseline adjustment. A microcontroller (Arduino Nano; Italy) acquires the analog phototransistor output at ∼1Hz and transmits the data to a computer running a Python-based interface for real-time visualisation and data logging. Complete electrical schematics details are provided in Figure S2.

#### Optical Signal Acquisition Principle (Beer-Lambert Law)

As blood perfusate flows through the transparent tubing, the system operates in transmission mode: the phototransistor output reflects the absorbance of the fluid based on Beer-Lambert principles (Eq. 1). Because ICG is administered as a discrete bolus and baseline absorbance is zeroed immediately prior to injection, subsequent changes in signal at 808 nm are attributed to changes in ICG concentration. Other chromophores, such as haemoglobin and bilirubin, are assumed to remain constant during the 90-minute clearance window. Notably, 808 nm is near an isosbestic point for oxy- and deoxyhaemoglobin (∼805 nm), minimising confounding from variations in oxygen saturation (Vos et al., 2014).

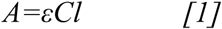

*Where A is absorbance, ε is the molar absorptivity, C is the concentration of absorbing species, and l is the optical path length*.

#### Optical Signal Acquisition Algorithm

To isolate ICG-specific absorbance signals and minimise noise, the sampling procedure included a method for removing background optical interference (Figure S3A). Under extreme ambient lighting conditions, corrected signals remained stable (Figure S3B).

### Optical Sensor Data Processing and Experimental Application

#### Test protocol

The sensor was clamped onto the circuit tubing and the photodiode sensitivity was adjusted so that measured pre-bolus absorbance was around 10% of full scale. After a 5-minute pre-bolus baseline to confirm signal stability, a 7.5 mg dose of indocyanine green (3 mL of 2.5 mg/mL) was injected directly into the turbulent suprahepatic inferior vena cava outflow of the perfusion reservoir to ensure optimal mixing, with the injection time logged on a custom GUI. Relative change in absorbance was continuously measured as an indicator of the rate of ICG clearance.

For each bolus, an analysis window was defined to represent the portion of the signal used for curve fitting and quantitative analysis. The start of the analysis window was empirically defined at t_0_ = 3 minutes, based on real-time optical traces demonstrating that all sensor signals had transitioned from the initial mixing phase to a stable clearance profile by this point. The analysis window extended up to 90 minutes post-bolus or until a stopping criterion was met. Three stopping criteria were applied: (i) the signal returned within 5 % of the pre-bolus baseline, (ii) interference occurred due to infusions or automated medications, or (iii) the sensor experienced a pronounced physical disturbance affecting the measured signal (Figure 4C). Additionally, a complete redo criterion was applied whereby (iv) the bolus was repeated if the signal undershot the pre-bolus baseline by more than 5 % of the baseline-to-peak amplitude range (observed in only 1 of the 45 clearance curves).

#### Sensor Linearisation and Pre-Processing Framework

A pre-processing framework was established by acquiring concurrent perfusate samples and analysing them via spectrophotometry (ThermoScientific Spectronic 200) to provide a ground-truth reference. The bolus and sampling protocol described by *Lau et al.* was adopted and performed in parallel with optical sensor recordings (Lau et al., 2024). Manual blood-perfusate samples were collected pre-bolus and then at 1, 3, 6, 9, 12, 15, 18, 21, 24, 30, 40, 50, 60, and 90 minutes post-bolus. Samples were drawn into serum separator tubes, centrifuged at 2800 RPM for 10 minutes, and the plasma absorbance measured at 805 nm and 950 nm with the final signal as 805 nm – 950 nm.

Continuous sensor data were paired with corresponding spectrophotometric readings to derive version-specific linearisation models (v1.0 = 5 boluses; v1.1 = 8 boluses). Hyperbolic functions were selected for both sensor versions as the optimal linearisation models, determined by the highest coefficients of determination (R^2^) amongst all paired datasets (Figure 4A). The arrangement of the sensor within the perfusion circuit and the associated data acquisition workflow are illustrated in Figure 2A.

#### Data Acquisition

Data acquisition followed a standardised five-step workflow to ensure reproducible ICG clearance measurements, as summarised in Figure 2B. Experimental procedures, including sensor placement, baseline acquisition, and bolus delivery, followed the test protocol. A custom program provided real-time signal visualisation, automated data logging at approximately 1 Hz, and event flagging of bolus injections. Continuous absorbance at 808 nm was recorded for up to 90 minutes post-injection or until an interfering event occurred.

#### Signal Pre-Processing

Following acquisition, raw absorbance traces were first linearised using the version-specific calibration functions derived in the Sensor Linearisation and Pre-Processing Framework section. The resulting signals were then normalised such that the 5-minute pre-bolus baseline corresponded to 0 and the start of the analysis period corresponded to 1 (arbitrary units). Subsequent analysis followed the standard test protocol.

#### Model and Quantify Clearance

Clearance behaviour was modelled using a bi-exponential decay function, an established approach for describing indocyanine green (ICG) elimination kinetics in the liver (Figure 3). The model is defined as:

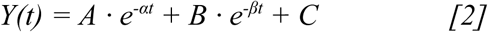

**Figure 3:**
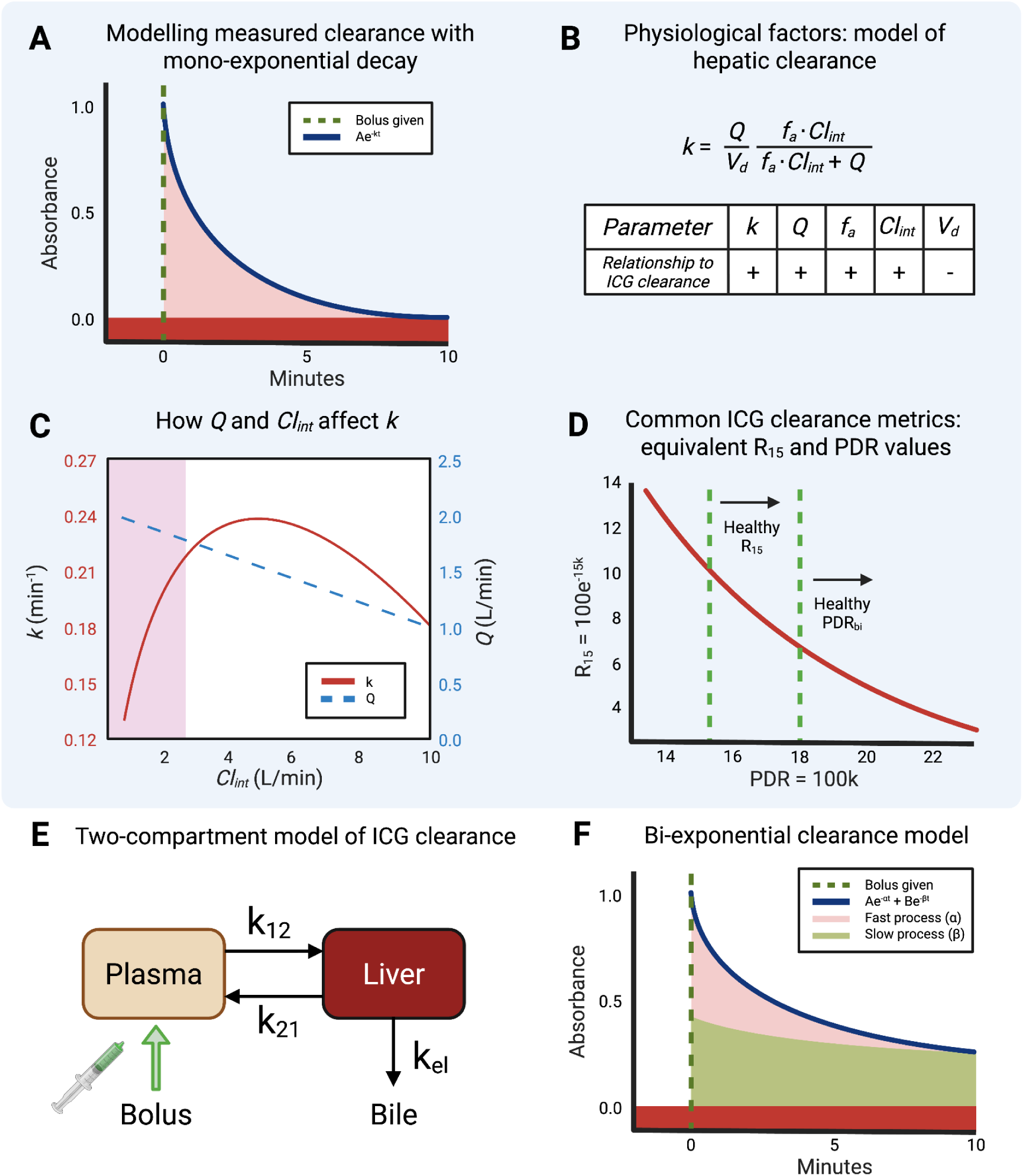
Modelling and interpreting ICG clearance curves. **(A)** The widely used mono-exponential fit (blue line) models ICG clearance as first-order exponential decay. In this model, parameter k is the elimination rate constant, representing the fraction of plasma concentration eliminated per unit time (Paumgartner et al., 1970). **(B)** The physiological determinants of k can be considered using an adapted hepatic clearance model (Köller et al., 2021), in which k is influenced by total hepatic blood flow (Q), the fraction of ICG available for uptake (f_a_), the intrinsic clearance of the hepatocytes (CL_int_) and the effective distribution volume (V_d_). **(C)** The physiological determinants of most interest are Q and Cl_int_ (Imamura et al., 2005). Examining the instantaneous k value (red line) for a given pair of Cl_int_ (x axis) and Q values (blue dashed line) reveals its sensitivity is range-dependent: when Cl_int_ is low it dominates k, even if Q is high (purple region). As Cl_int_ increases, k becomes limited by Q (white region). **(D)** Two established metrics for ICG clearance (both dependent solely on k) are R15, the percentage of ICG remaining after 15 minutes, and PDR, the percentage of ICG cleared per minute. These metrics are related but do not vary linearly with each other, and there is some variation in established thresholds for abnormal clearance (Vos et al., 2014). **(E)** The two-compartment model is a more detailed representation of ICG clearance, distinguishing rapid uptake of ICG from plasma into hepatocytes (k_12_, k_12_) from its subsequent hepatobiliary elimination (k_el_) (Imamura et al., 2005). **(F)** Solving the two-compartment model yields a bi-exponential decay with two hybrid rate constants: a fast phase (α), reflecting initial hepatocyte uptake, and a slow phase (β), reflecting hepatobiliary excretion (Gasperi et al., 2016). Beyond capturing potential information on uptake, this model robustly fits ICG clearance curves that deviate from mono-exponential decay, making it a superior general-purpose model for ICG clearance. Created in BioRender. Derwent, E. (2025) https://BioRender.com/d93gmzn

where C represents the fixed baseline, defined as the mean of the 5-minute pre-bolus window. Curve fitting was performed using nonlinear least-squares optimisation, and only data within the defined analysis window were included. Model performance was quantified by the coefficient of determination (R^2^).

From the fitted curve, two clearance metrics were derived:

**15-minute retention (*R_15_*, %):** calculated analytically from the fitted function as the proportion of the initial signal (at t_0_) remaining 15 minutes after the t_0_ value:

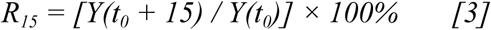

**Plasma disappearance rate (PDR_bi_, %/min):** obtained by numerically solving the fitted bi-exponential equation for the time at which the signal decayed to half of its initial peak (PD_50_)

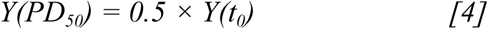

PDR_bi_ was then calculated as:

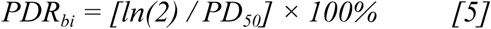

R_15_ represents the percentage of ICG retained 15 minutes after t₀, whereas PDR_bi_ expresses the overall rate of plasma clearance as a percentage per minute. Both metrics are standard endpoints in clinical ICG testing and serve as established indicators of hepatocellular clearance capacity (Gasperi et al., 2016; Vos et al., 2014; Wagener, 2013). Together, these parameters quantified ICG clearance kinetics and enabled comparison of liver performance and health across experimental conditions.

## Results

### Sensor Validation and Signal Characteristics

Across 13 human livers (10 whole, 3 split), we recorded 45 ICG clearance curves, including 14 paired with spectrophotometric sampling for linearisation. Linearised sensor output correlated closely with spectrophotometric absorbance values (see Figure 4A), yielding a mean coefficient of determination (R^2^) of 0.996 (range 0.983 - 0.999). Each sensor version maintained effective calibration performance on all clearance curves, with v1.0 achieving an R^2^ range of 0.990 - 0.999 and v1.1 achieving 0.983 - 0.999.

**Figure 4:**
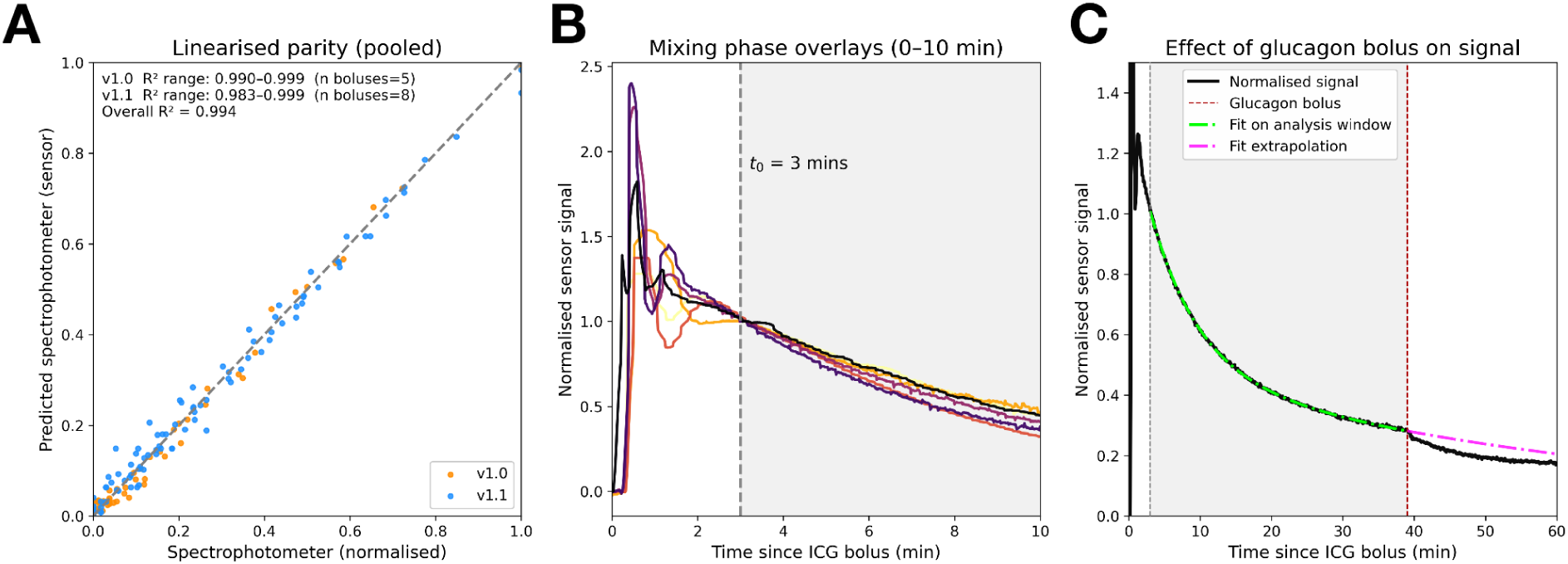
Linearisation accuracy, mixing-phase behaviour, and exclusion of pharmacologic interference. **(A)** Linearised sensor signal agrees with spectrophotometer across boluses (pooled R^2^ = 0.994); v1.0 range 0.990–0.999 (n=5) and v1.1 range 0.983–0.999 (n=8), confirming the per-version calibration. **(B)** 6 Exemplar overlays (0 → 10 min) show the mixing phase finishing by 3 min (dashed line); beyond this point traces follow a smooth decay suitable for fitting. **(C)** A glucagon bolus (red dashed line) perturbs the signal outside the analysis window (grey). The model is fit only within the window (green) and shown extrapolated (magenta) to illustrate why the affected segments are excluded from the curve fit.

Continuous optical signals exhibited a short mixing phase (<3 min) before reaching a stable clearance profile (Figure 4B). Beyond this point, all boluses followed a smooth mono- or bi-exponential decay curve, characteristic of hepatic elimination. Signal artefacts from infusions or physical disturbances were infrequent and readily explainable (e.g. glucagon bolus; Figure 4C). The resulting analysis window isolated the physiologic clearance phase, ensuring that fitted curves best represented true ICG elimination dynamics.

### ICG Clearance Across Varying Temperature Conditions

ICG clearance kinetics were consistently faster during NMP compared with SNMP (Figure 5A–5D). Across all four livers (W1, W2, W3, W4), the mean increase in PDR_bi_ (ΔPDR_bi_) was +14.41 %/min (8.15 vs. 22.56 %/min; range: 3.07 - 24.24%/min), while the 15-minute retention (R_15_) decreased by 26.2 percentage points (32.0% vs. 9.65%; range: –12.6 - –30.9 percentage points) for NMP and SNMP, respectively. Each NMP trace exhibited a steeper exponential decay and lower residual signal relative to its paired SNMP curve. In a separate experiment on a single liver, clearance was faster at 36 °C than at 32 °C, with ΔPDR_bi_ = +2.57 %/min (9.31 vs. 11.88 %/min) and ΔR15 = −6.0 percentage points (33.9 % vs. 27.9 %), respectively. Collectively, these results demonstrate that normothermic temperature (35.5°C to 37.5°C) (Van Beekum et al., 2021) produces higher ICG clearance rates independent of perfusate composition.

**Figure 5:**
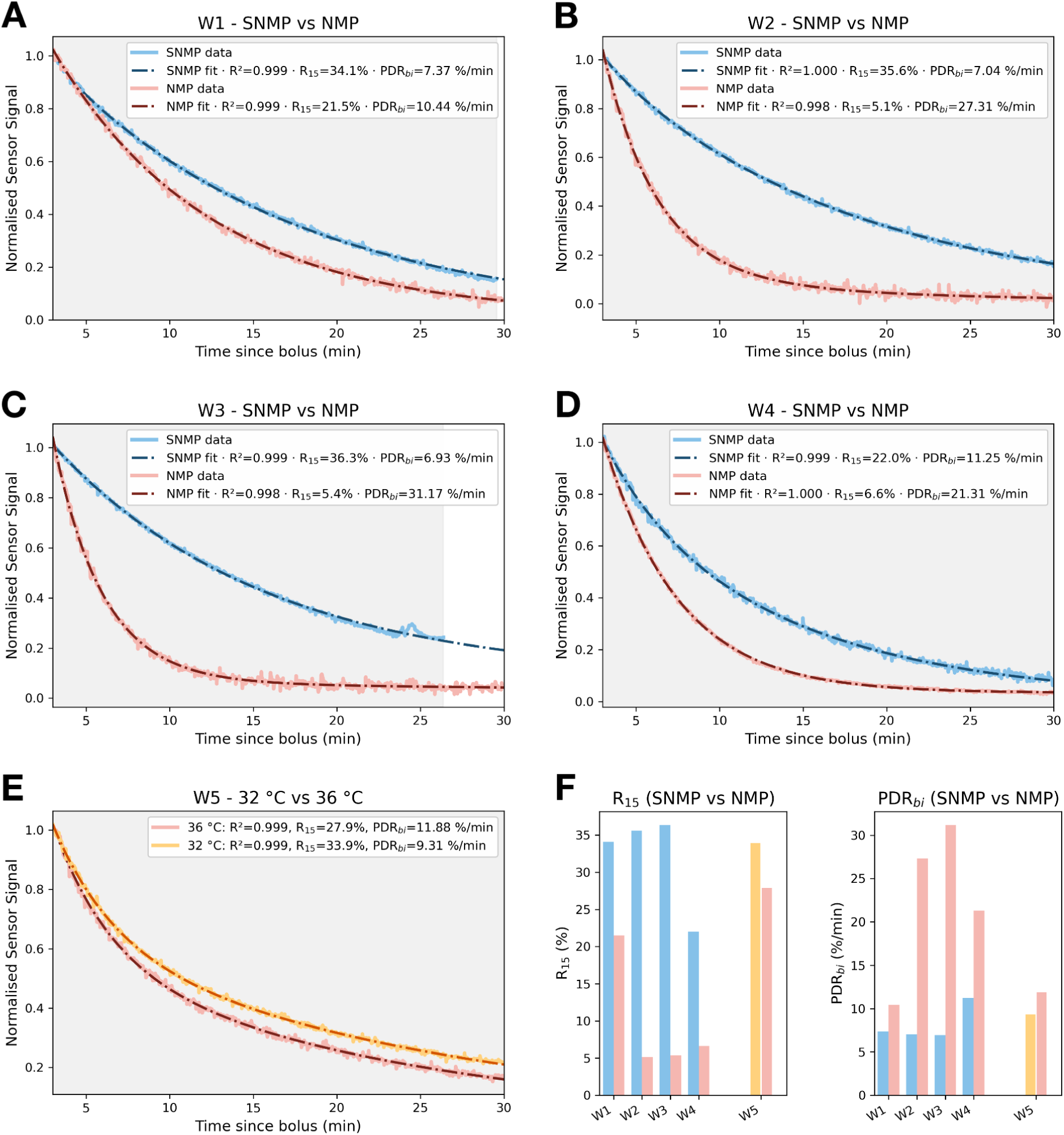
Indocyanine green (ICG) clearance under SNMP and NMP conditions at 32 °C and 36 °C. Representative clearance curves for livers W1–W5 **(A–E)** show normalised sensor absorbance signals (solid lines) with corresponding bi-exponential fits (dashed lines). Each fit reports coefficient of determination (R^2^), 15-minute retention (R_15_), and biexponential plasma disappearance rate (PDR_bi_), calculated from the curve based on analysis period (shaded background). **(A-D)** Across all livers (W1–W4), normothermic machine perfusion (NMP, red) exhibited faster clearance kinetics (lower R_15_, higher PDR_bi_) than subnormothermic perfusion (SNMP, blue). **(E)** Comparison of ICG boluses at 32 °C and 36 °C (W5) shows that clearance is faster at normothermic (36 °C) temperatures (R_15_: 33.9 → 27.9%, PDR: 9.31 → 11.88%). **(F)** Paired column plots summarise R_15_ and PDR_bi_ for each liver, highlighting the consistent improvement in clearance during NMP.

### Model Performance and Clearance Metric Behaviour

Bi-exponential fitting achieved substantially higher agreement with measured ICG clearance data than mono-exponential fitting (Figure 6). Across all 45 boluses, mono-exponential fits produced R^2^ values between 0.849 and 0.999 (median = 0.982), whereas bi-exponential fits achieved R^2^ values between 0.965 and 0.9997 (median = 0.999) (Figure S4). In several cases, particularly those with slow or biphasic elimination, mono-exponential models substantially underperformed (ΔR^2^ ≤ 0.15), producing distorted R_15_ and PDR estimates relative to bi-exponential fits. Four representative examples are shown in Figure 6, illustrating the systematic underestimation of clearance speed (i.e., 68.5 % vs. 39.1 % R_15_; 2.52 %/min vs. 7.42 %/min). The in-silico simulation of 10,000 bi-exponential decays further demonstrated that R_15_ and PDR_bi_ provide robust, interpretable metrics that capture physiologically plausible clearance behaviour across a wide range of kinetic profiles.

**Figure 6:**
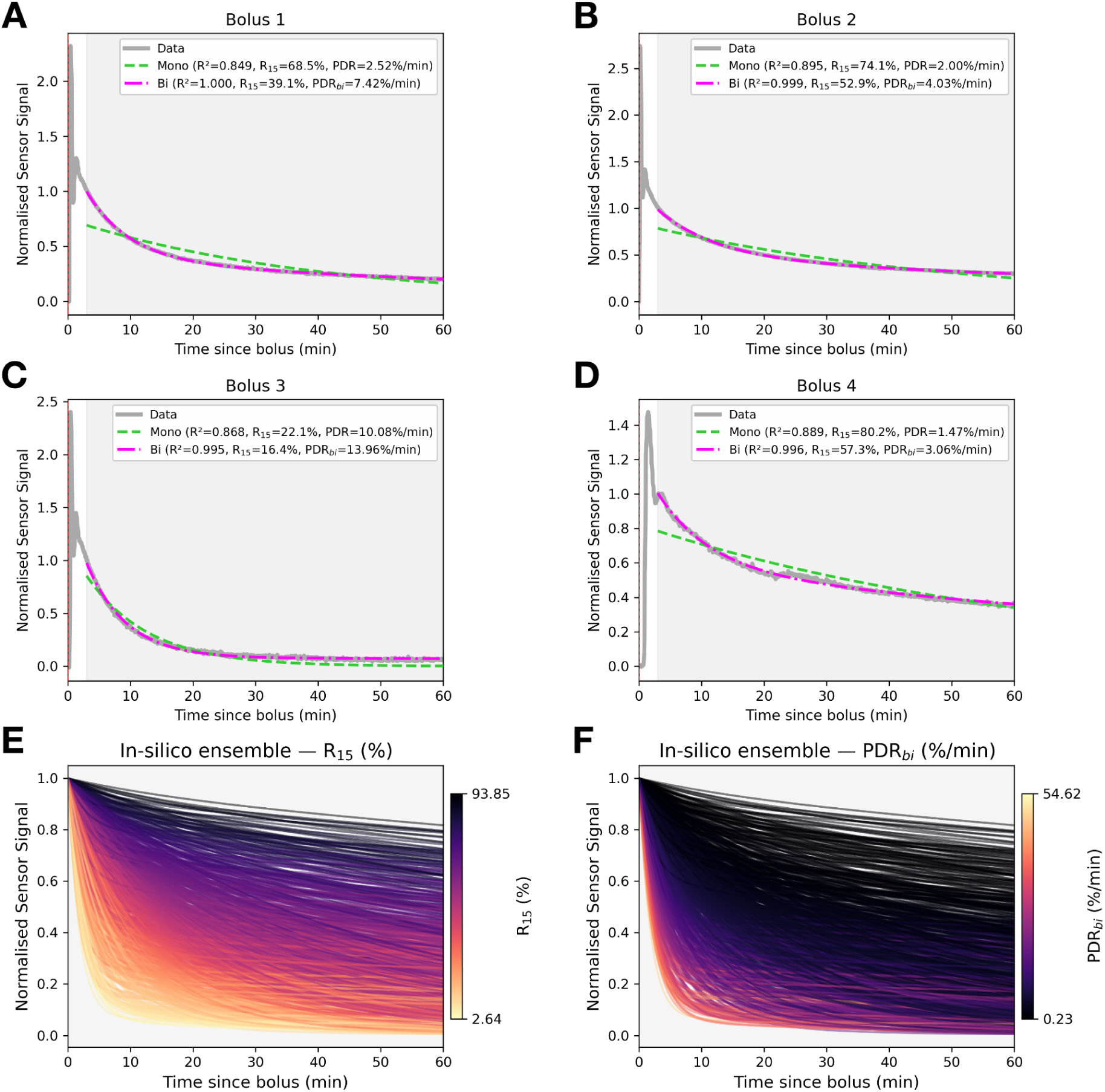
Comparison of mono-exponential and bi-exponential fitting models for ICG clearance, with in-silico simulations illustrating behaviour of clearance metrics. **(A-D)** Example ICG clearance curves from four boluses demonstrate that mono-exponential models (cyan) can underestimate clearance kinetics during machine perfusion, whereas bi-exponential fits (magenta) provide superior agreement with measured data (Mono-exponential R^2^: 0.849-0.999; R^2^_median_: 0.982 vs Bi-exponential R²: 0.965-0.9997; R^2^_median_: 0.999). Each fit reports R_15_ and PDR_bi_. **(E-F)** show ensembles of 10,000 simulated bi-exponential decays coloured by (E) R_15_ and (F) PDR_bi_, confirming that these metrics accurately track overall clearance rate across variation in bi-exponential decay curves.

## Discussion

This study presents the first demonstration of real-time optical measurement of ICG clearance in an ex-vivo liver perfusion system. The device is a low-cost, reusable, clamp-on sensor that enables continuous, non-invasive assessment of hepatic function through uninterrupted optical acquisition of the clearance profile. The sensor removes the need for repeated manual sampling and associated handling time, while maintaining high agreement with ground-truth spectrophotometry measurements. Application of the sensor across perfusion conditions revealed consistently faster ICG elimination during NMP relative to SNMP and 32 °C perfusion, consistent with elevated metabolic activity at physiological temperatures and reinforcing ICG clearance as a surrogate marker of metabolic function. To our knowledge, this represents the first application of ICG clearance measurement under subnormothermic, acellular perfusion conditions.

Constructed from off-the-shelf components and deployable without need for sterilisation, the sensor is transferable across perfusion platforms and enables standardised, dynamic graft assessment. Optical systems such as the LiMON device have previously enabled non-invasive, transcutaneous ICG clearance monitoring in vivo (De Liguori Carino et al., 2009); however, no equivalent technique has been applied to ex-vivo perfusion, where continuous, platform-independent monitoring can provide direct insight into graft metabolic performance (Gasperi et al., 2016; Levesque et al., 2016; Vos et al., 2014).

Continuous optical monitoring revealed a short, high-variance mixing phase immediately following ICG administration, lasting up to three minutes before stabilisation of the clearance curve. This period, dominated by convective and diffusive mixing artefacts, could not be resolved using conventional discrete sampling protocols that typically obtain the first measurements within this mixing phase (Gasperi et al., 2016; Lau et al., 2024; Levesque et al., 2016). As a result, early concentration readings in previous studies likely reflected non-homogeneous perfusate distribution rather than true hepatic clearance. The sensor also detected transient perturbations in the clearance signal associated with medication administration, such as glucagon boluses - which likely would have been overlooked by discrete sampling - allowing these events to be identified and excluded from curve fitting.

The optical sensor achieved measurement accuracy comparable to the spectrophotometric ground truth while providing substantially higher temporal resolution. Linearisation of the raw absorbance data yielded highly consistent signals, allowing ICG clearance to be modelled with precision using the sensor. Comparison with discrete spectrophotometry showed near-perfect correlation, confirming the validity of the optical method for continuous use. Across the full dataset, bi-exponential fitting achieved uniformly high agreement with measured signals, while mono-exponential fits showed greater variability and performed adequately only in a subset of traces with simple decay behaviour. These findings demonstrate that the optical system reproduces spectrophotometric accuracy while enabling quantitative modelling of ICG clearance with greater fidelity and much greater temporal definition than discrete sampling methods (Lau et al., 2024; Levesque et al., 2016; Vos et al., 2014).

While mono-exponential fitting offers simplicity and has been widely adopted in clinical ICG clearance analysis (Gasperi et al., 2016; Lau et al., 2024; Levesque et al., 2016), our data show that this approach fails to capture the kinetics of many perfused livers, particularly those exhibiting slower or biphasic elimination. In such cases, the mono-exponential model consistently underrepresented R_15_ and PDR estimates, distorting true hepatic clearance behaviour. The bi-exponential formulation overcomes these limitations by independently resolving the rapid distribution and slower elimination phases (Figure 3F), providing a more physiologically consistent quantification of ICG clearance (Imamura et al., 2005; Vos et al., 2014). Across all boluses, this model consistently reproduced measured curves with equivalent or higher R^2^ and enabled reliable extraction of both R_15_ and PDR_bi_ parameters. Based on the results presented, the bi-exponential approach represents a more physiologically accurate representation of ICG clearance.

Because the bi-exponential model decomposes ICG elimination into two distinct kinetic components - and therefore two decay constants (ɑ, β) - traditional mono-exponential metrics such as half-life and PDR cannot be directly calculated from the fitted curve (Gasperi et al., 2016; Ge et al., 2014; Lau et al., 2024). Instead, calculations for clearance metrics R_15_ and PDR_bi_ based on the bi-exponential model were proposed, to preserve existing intuition and decision-making frameworks regarding hepatic function. Experimental results showed that both parameters reflected differences in metabolic activity, with higher PDR_bi_ and lower R_15_ observed under NMP compared with SNMP conditions. This temperature-dependent reduction in clearance rate mirrors the diminished ICG elimination observed *in vivo* in patients with impaired hepatic metabolism (De Liguori Carino et al., 2009; Ohwada et al., 2006). Clinically, ICG clearance also distinguishes recovery and dysfunction: Merle *et al*. reported significantly higher PDRs in patients who recovered from acute liver failure compared with those who did not, while Schwarz *et al*. demonstrated that preoperative ICG clearance strongly predicted postoperative hepatic dysfunction (Gasperi et al., 2016; Levesque et al., 2016; Merle et al., 2009; Schwarz et al., 2018). While these metrics inevitably condense a complex, multiphasic process into single values, they remain clinically meaningful proxies for liver function, providing continuity with existing literature and enabling comparison and understanding of liver function across both clinical and ex-vivo contexts.

Several limitations of the current sensor system should be acknowledged. From a hardware perspective, we currently employ an absorbance (transmission) configuration; future iterations might benefit from exploring a reflectance setup, which is well described in sensor literature as feasible alternatives to combat optical coupling issues (O’Toole and Diamond, 2008). The sensor currently measures absorbance through whole blood rather than serum, introducing potential spectral interference from hemoglobin and other chromophores. However, this effect is likely minimal, as baseline drift during clearance recordings was negligible across most boluses and the peak absorbance wavelength of ICG is the isosbestic point of oxyHb and deoxyHb, meaning changes in oxygenation should negligibly affect readings (Vos et al., 2014). Incorporating an auxiliary wavelength, such as 950 nm as used by Lau *et al*. or 905 nm proposed by Vos *et al*., would allow real-time tracking of baseline stability and correction for slow optical shifts (Lau et al., 2024; Vos et al., 2014). Nevertheless, the linearised sensor output demonstrated very strong agreement with spectrophotometer readings corrected with 950 nm reference data. As a dynamic perfusion system, clearance signals can also be transiently affected by pharmacologic or mechanical events, such as glucagon boluses or pump adjustments, although these are readily identifiable in continuous recordings. Finally, this work remains limited to ex-vivo experiments; the absence of clinical outcome data precludes correlation of sensor-derived metrics with transplant viability or postoperative function. Future validation in clinical and translational settings will be required to establish predictive relevance of the sensor and metrics.

Several avenues could extend this work. From an engineering standpoint, future iterations should focus on electrical and optical optimisation, including improved signal conditioning, integrated baseline detection, and automated bolus flagging and artefact identification. Hardware development will also explore reflectance configurations to improve long-term optical stability. These refinements aim to increase signal robustness, reduce alignment sensitivity, and render laborious manual intervention virtually obsolete, moving the system toward fully autonomous operation across perfusion platforms.

Beyond improvements to the sensor itself, the ex-vivo perfusion context provides a uniquely controllable environment for mechanistic study. Because flow rate, perfusate composition, and bile production (among many other parameters) can be precisely measured or even independently adjusted, the system enables quantitative testing of pharmacokinetic clearance models under well-defined physiological conditions. In contrast, in vivo assessment of ICG clearance is inherently confounded by the inability to isolate or quantify hepatic blood flow, circulating volume, biliary output and intrinsic hepatic clearance - factors that jointly influence ICG kinetics through their effects on perfusion, hepatocellular uptake, and canalicular excretion (Levesque et al., 2016; Sakka, 2018). By fixing or directly measuring these variables, ex-vivo perfusion can isolate the hepatic contribution to ICG uptake and excretion, providing a controlled platform to interrogate the mechanisms governing dye extraction and elimination. This setting enables exploration of distinct kinetic models, such as the bi-exponential two-component model - (1) representing rapid vascular uptake and (2) slower biliary excretion - first proposed by Paumgartner *et al*. and corroborated by Köller *et al*. (Köller et al., 2021; Paumgartner et al., 1970). It also facilitates more advanced analyses, including the ratio of fast-to-slow clearance components, the impact of perfusate bilirubin or albumin content, and the direct correlation of slow-phase kinetics with biliary excretion. Future work could extend this approach further by integrating a sensor on the bile-output tubing, enabling real-time correlation of ICG clearance with biliary output as bile production and hepatic function are further optimised.

The sensor should next be implemented in a prospective clinical trial to validate the bi-exponential model and associated clearance metrics against patient outcomes. This will determine which parameters - R_15_, PDR_bi_, or phase-specific decay rates - best predict post-transplant graft function and long term patient outcomes. In parallel, the broader platform is adaptable to other optically traceable clearance probes where the liver is primarily responsible for elimination or metabolism. Optically active analytes with defined absorbance spectra, such as caffeine (absorbance peak λ ≈ 274 nm) (Ahmad Bhawani et al., 2015), may be targeted using analogous non-invasive optical sensing architectures for quantitative evaluation of clearance kinetics. Extending the same experimental advantages demonstrated with ICG, the controlled nature of machine perfusion permits equivalent clearance studies for these other optically active compounds, where perfusion variables can be tightly controlled and system behaviour continuously monitored. This positions the sensor-perfusion platform as a versatile tool for comparative clearance analysis across an array of targets.

## Conclusion

This study demonstrates that continuous, real-time optical monitoring of ICG clearance during ex-vivo liver perfusion provides an accurate and more practical method for assessing hepatic function compared to previous sampling-based techniques. The clamp-on optical sensor achieved near-perfect agreement with spectrophotometric ground truth while enabling high-resolution kinetic modelling unobtainable by intermittent sampling. Using this approach, ICG clearance was shown to increase with metabolic activity between NMP and SNMP conditions, and the derived metrics R_15_ and PDR captured these physiological differences with high fidelity. Together, these findings establish non-invasive optical ICG monitoring during ex-vivo perfusion as a robust quantitative tool for dynamic graft assessment and for advancing mechanistic understanding of hepatic clearance. Future work will focus on clinical validation of these metrics in transplant settings and extension of the sensing platform to additional optically active analytes, broadening its utility as a versatile framework for real-time clearance analysis across perfusion and physiological systems.

## Financial support and sponsorship

None

## Conflicts of interest

Nothing to report

## Acknowledgements

The authors would like to acknowledge the support of COARO and RPATI

## Author contributions

**Concept/design:** Derwent ENJ, Babekuhl D

**Data collection:** Derwent ENJ, Babekuhl D, Risbey CWG, Niu A, Yousif P

**Data analysis/interpretation:** Derwent ENJ, Babekuhl D, Risbey CWG

**Perfusion systems engineering:** Babekuhl D, Fonseka N, Derwent ENJ, Curry S, Seow C, Ng I

**Drafting article:** Derwent ENJ, Babekuhl D, Risbey CWG

**Critical revision of article:** Derwent ENJ, Babekuhl D, Risbey CWG, Niu A, Fonseka N, Curry S, Seow C, Crawford M, Pulitano C

**Approval of article:** All authors

**Funding secured by:** Pulitano C, McCaughan GW, Crawford M

## Declaration of generative AI and AI-assisted technologies in the manuscript preparation process

During the preparation of this work, the authors used ChatGPT (Open AI) to:

1. Refine language, grammar, and phrasing across the manuscript and
2. Assist with Python coding for the linearisation and the data analysis pipeline.

After using this tool, the authors reviewed, edited, tested, and validated all text and code as needed and take full responsibility for the content of the published article. No generative AI systems are listed as authors, and no AI tools were used to generate data, images or conclusions.

## Notes

### Competing Interest Statement

The authors have declared no competing interest.

## References

Ahmad Bhawani, S., Fong, S.S., Mohamad Ibrahim, M.N., 2015. Spectrophotometric Analysis of Caffeine. Int. J. Anal. Chem. 2015, 1–7. 10.1155/2015/170239

Alam, K., Crowe, A., Wang, X., Zhang, P., Ding, K., Li, L., Yue, W., 2018. Regulation of Organic Anion Transporting Polypeptides (OATP) 1B1- and OATP1B3-Mediated Transport: An Updated Review in the Context of OATP-Mediated Drug-Drug Interactions. Int. J. Mol. Sci. 19, 855. 10.3390/ijms19030855

Babekuhl, D., Risbey, C., Niu, A., Yousif, P., Liu, K., McCaughan, G., Strasser, S., Crawford, M., Pulitano, C., 2025. Normothermic Preservation of Human Livers for More Than 3 Weeks: From Experimental Model to Clinical Application. Am. J. Transplant. 25, S95. 10.1016/j.ajt.2025.07.198

Brüggenwirth, I.M.A., De Meijer, V.E., Porte, R.J., Martins, P.N., 2020. Viability criteria assessment during liver machine perfusion. Nat. Biotechnol. 38, 1260–1262. 10.1038/s41587-020-0720-z

De Liguori Carino, N., O’Reilly, D.A., Dajani, K., Ghaneh, P., Poston, G.J., Wu, A.V., 2009. Perioperative use of the LiMON method of indocyanine green elimination measurement for the prediction and early detection of post-hepatectomy liver failure. Eur. J. Surg. Oncol. EJSO 35, 957–962. 10.1016/j.ejso.2009.02.003

Destro, M., Puliafito, C.A., 1989. Indocyanine Green Videoangiography of Choroidal Neovascularization. Ophthalmology 96, 846–853. 10.1016/S0161-6420(89)32826-0

Devarbhavi, H., Asrani, S.K., Arab, J.P., Nartey, Y.A., Pose, E., Kamath, P.S., 2023. Global burden of liver disease: 2023 update. J. Hepatol. 79, 516–537. 10.1016/j.jhep.2023.03.017

Fan, Z., Zong, J., Lau, W.Y., Zhang, Y., 2020. Indocyanine green and its nanosynthetic particles for the diagnosis and treatment of hepatocellular carcinoma. Am. J. Transl. Res. 12, 2344–2352.

Ferrigno, A., 2015. Metabolic shift in liver: Correlation between perfusion temperature and hypoxia inducible factor-1α. World J. Gastroenterol. 21, 1108. 10.3748/wjg.v21.i4.1108

Gasperi, A.D., Mazza, E., Prosperi, M., 2016. Indocyanine green kinetics to assess liver function: Ready for a clinical dynamic assessment in major liver surgery? World J. Hepatol. 8, 355. 10.4254/wjh.v8.i7.355

Ge, P.-L., Du, S.-D., Mao, Y.-L., 2014. Advances in preoperative assessment of liver function. Hepatobiliary Pancreat. Dis. Int. 13, 361–370. 10.1016/S1499-3872(14)60267-8

Girgenti, R., Tropea, A., Buttafarro, M.A., Ragusa, R., Ammirata, M., 2020. Quality of Life in Liver Transplant Recipients: A Retrospective Study. Int. J. Environ. Res. Public. Health 17, 3809. 10.3390/ijerph17113809

Imamura, H., Sano, K., Sugawara, Y., Kokudo, N., Makuuchi, M., 2005. Assessment of hepatic reserve for indication of hepatic resection: decision tree incorporating indocyanine green test. J. Hepatobiliary. Pancreat. Surg. 12, 16–22. 10.1007/s00534-004-0965-9

Kalliokoski, A., Niemi, M., 2009. Impact of OATP transporters on pharmacokinetics. Br. J. Pharmacol. 158, 693–705. 10.1111/j.1476-5381.2009.00430.x

Köller, A., Grzegorzewski, J., König, M., 2021. Physiologically Based Modeling of the Effect of Physiological and Anthropometric Variability on Indocyanine Green Based Liver Function Tests. Front. Physiol. 12, 757293. 10.3389/fphys.2021.757293

Lau, N., Ly, M., Ewenson, K., Toomath, S., Ly, H., Mestrovic, N., Liu, K., McCaughan, G., Crawford, M., Pulitano, C., 2024. Indocyanine green: A novel marker for assessment of graft quality during ex situ normothermic machine perfusion of human livers. Artif. Organs 48, 472–483. 10.1111/aor.14696

Levesque, E., Martin, E., Dudau, D., Lim, C., Dhonneur, G., Azoulay, D., 2016. Current use and perspective of indocyanine green clearance in liver diseases. Anaesth. Crit. Care Pain Med. 35, 49–57. 10.1016/j.accpm.2015.06.006

Merle, U., Sieg, O., Stremmel, W., Encke, J., Eisenbach, C., 2009. Sensitivity and specificity of plasma disappearance rate of indocyanine green as a prognostic indicator in acute liver failure. BMC Gastroenterol. 9, 91. 10.1186/1471-230X-9-91

Noitumyae, J., 2021. Indocyanin greens cholangiography for intra-operative bile duct visualization during pediatric laparoscopic hepato-biliary surgery. J. Pediatr. Surg. Case Rep. 69, 101833. 10.1016/j.epsc.2021.101833

Ohwada, S., Kawate, S., Hamada, K., Yamada, T., Sunose, Y., Tsutsumi, H., Tago, K., Okabe, T., 2006. Perioperative real-time monitoring of indocyanine green clearance by pulse spectrophotometry predicts remnant liver functional reserve in resection of hepatocellular carcinoma. Br. J. Surg. 93, 339–346. 10.1002/bjs.5258

O’Toole, M., Diamond, D., 2008. Absorbance Based Light Emitting Diode Optical Sensors and Sensing Devices. Sensors 8, 2453–2479. 10.3390/s8042453

Paumgartner, G., Probst, P., Kraines, R., Leevy, C.M., 1970. KINETICS OF INDOCYANINE GREEN REMOVAL FROM THE BLOOD*. Ann. N. Y. Acad. Sci. 170, 134–147. 10.1111/j.1749-6632.1970.tb37009.x

Risbey, C.W.G., Babekuhl, D., Yousif, P., Fonseka, N., Zhang, W.B., Derwent, E., Curry, S., Seow, C., Niu, A., Liu, K., Strasser, S.I., McCaughan, G.W., Crawford, M., Pulitano, C., 2025. A novel, institutionally developed hypothermic oxygenated machine PErfusion system allows low-cost, universal implementation for liver transplantation: A safety and feasibility pilot study. Liver Transpl. 10.1097/LVT.0000000000000686

Risbey, C.W.G., Lau, N.-S., Niu, A., Zhang, W.B., Crawford, M., Pulitano, C., 2024. Return of the cold: How hypothermic oxygenated machine perfusion is changing liver transplantation. Transplant. Rev. 38, 100853. 10.1016/j.trre.2024.100853

Sakka, S.G., 2018. Assessment of liver perfusion and function by indocyanine green in the perioperative setting and in critically ill patients. J. Clin. Monit. Comput. 32, 787–796. 10.1007/s10877-017-0073-4

Schlegel, A., Muller, X., Dutkowski, P., 2019. Machine perfusion strategies in liver transplantation. HepatoBiliary Surg. Nutr. 8, 490–501. 10.21037/hbsn.2019.04.04

Schurink, I.J., Luijmes, S.H., Willemse, J., De Goeij, F.H.C., Groen, P.C., Küçükerbil, E.H., Broere, R., Pascale, M.M., Porte, R.J., Tintu, A.N., Van Der Laan, L.J.W., Polak, W.G., De Jonge, J., 2025. Assessment of Ex Situ Liver Function by Indocyanine Green Clearance During Clinical Normothermic Machine Perfusion of Extended Criteria Grafts. Transplantation 109, e484–e493. 10.1097/TP.0000000000005350

Schwarz, C.C., Plass, I., Fitschek, F., Mittlböck, M., Kampf, S., Asenbaum, U., Starlinger, P., Stremitzer, S., Bodingbauer, M., Kaczirek, K., 2018. Value of Indocyanine Green Clearance Assessment to Predict Postoperative Liver Dysfunction in Patients Undergoing Liver Resection. J. Am. Coll. Surg. 227, S183. 10.1016/j.jamcollsurg.2018.07.366

Van Beekum, C.J., Vilz, T.O., Glowka, T.R., Von Websky, M.W., Kalff, J.C., Manekeller, S., 2021. Normothermic Machine Perfusion (NMP) of the Liver – Current Status and Future Perspectives. Ann. Transplant. 26. 10.12659/AOT.931664

Vos, J.J., Wietasch, J.K.G., Absalom, A.R., Hendriks, H.G.D., Scheeren, T.W.L., 2014. Green light for liver function monitoring using indocyanine green? An overview of current clinical applications. Anaesthesia 69, 1364–1376. 10.1111/anae.12755

Wagener, G., 2013. Assessment of Hepatic Function, Operative Candidacy, and Medical Management after Liver Resection in the Patient with Underlying Liver Disease. Semin. Liver Dis. 33, 204–212. 10.1055/s-0033-1351777

Wheeler, H.O., Cranston, W.I., Meltzer, J.I., 1958. Hepatic Uptake and Biliary Excretion of Indocyanine Green in the Dog. Exp. Biol. Med. 99, 11–14. 10.3181/00379727-99-24229

