## Supplementary material for "Development and Validation of a Continuous Real-Time Optical Sensor for Indocyanine Green Clearance Measurement During Ex-Vivo Perfusion of Human Livers": Figure S1

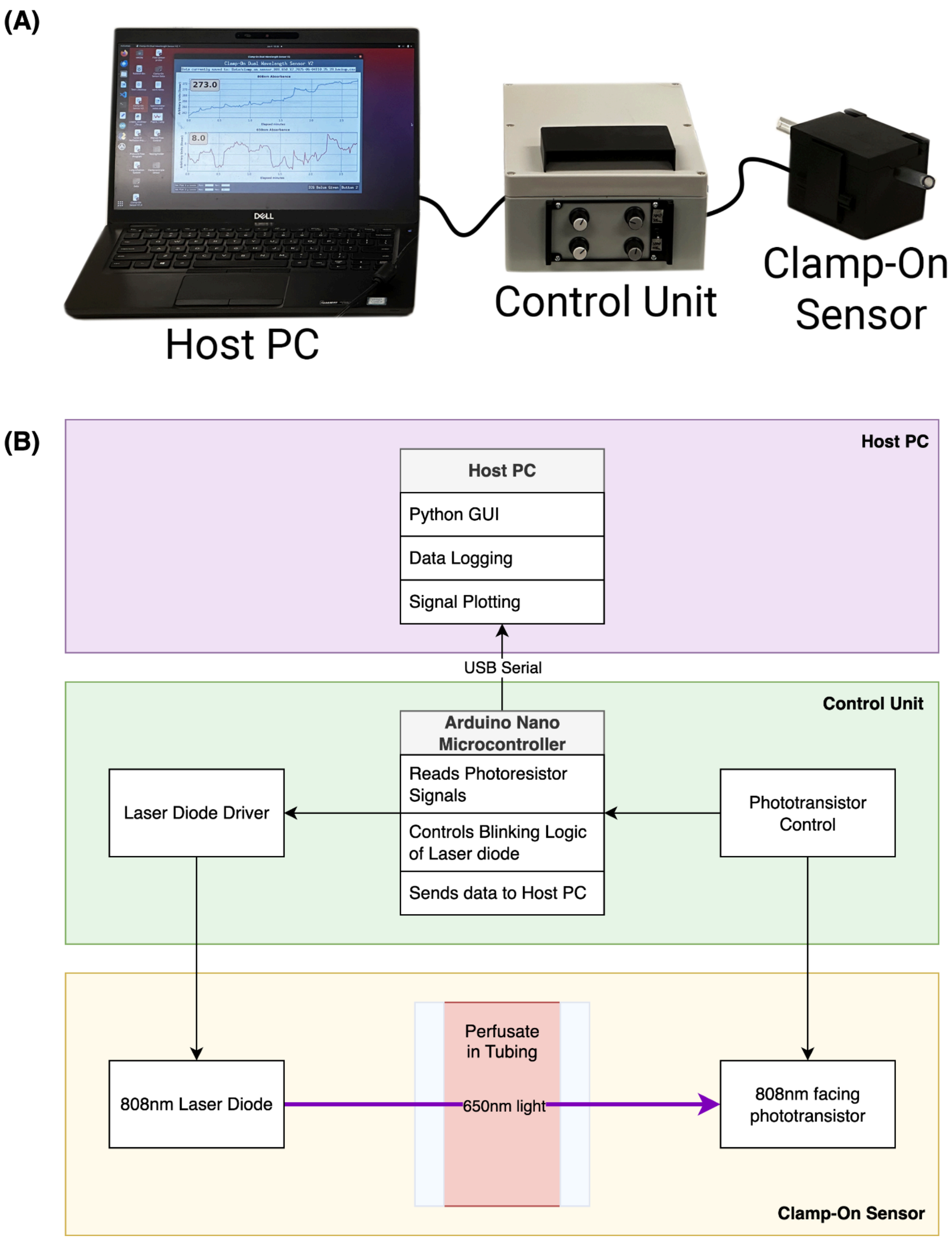


***Figure S1: (A) Physical system overview.*** *Left: Data acquisition and visualisation PC running custom Python software. Centre: Electronics unit containing laser drivers and phototransistor potentiometer control. Right: Clamp-on optical sensor positioned on sample perfusion tubing (Tygon ND-100-65, ¼" ID, ⅜" OD).* ***(B) Block diagram illustrating the system architecture.*** *The host PC runs a custom Python interface for data visualisation, logging, and control via USB*

*serial communication with the control unit. The control unit, based on an Arduino Nano microcontroller, manages laser diode operation, reads phototransistor signals, and regulates detector sensitivity. The clamp-on sensor houses an 808 nm laser diode and opposing phototransistor positioned across transparent perfusion tubing to measure transmitted light intensity and derive relative absorbance.*
