## Supplementary material for "Development and Validation of a Continuous Real-Time Optical Sensor for Indocyanine Green Clearance Measurement During Ex-Vivo Perfusion of Human Livers": Figure S2

(A)

### V1.0 Sensor Schematic

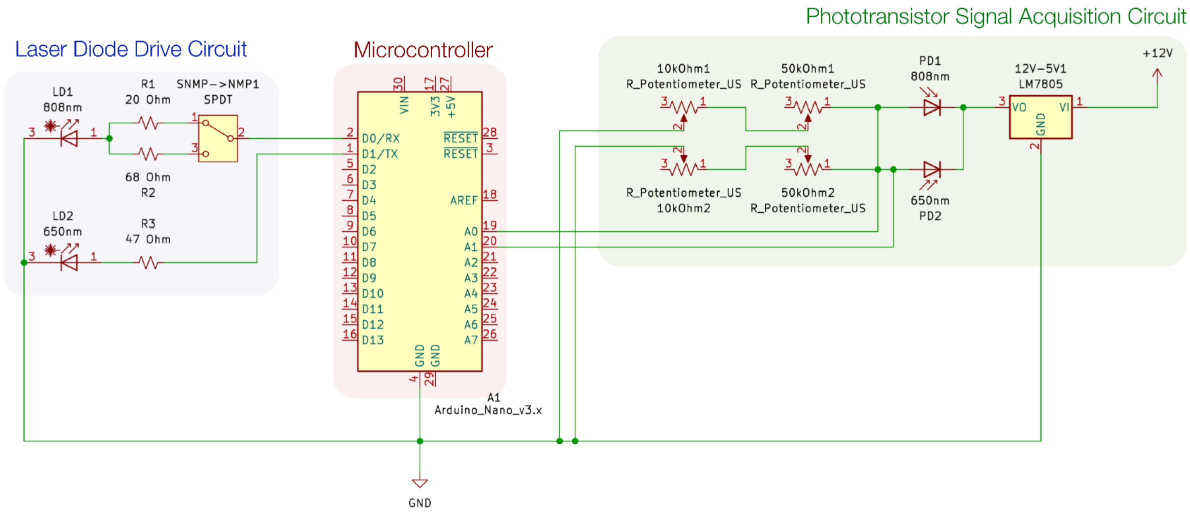


(B)

### V1.1 Sensor Schematic

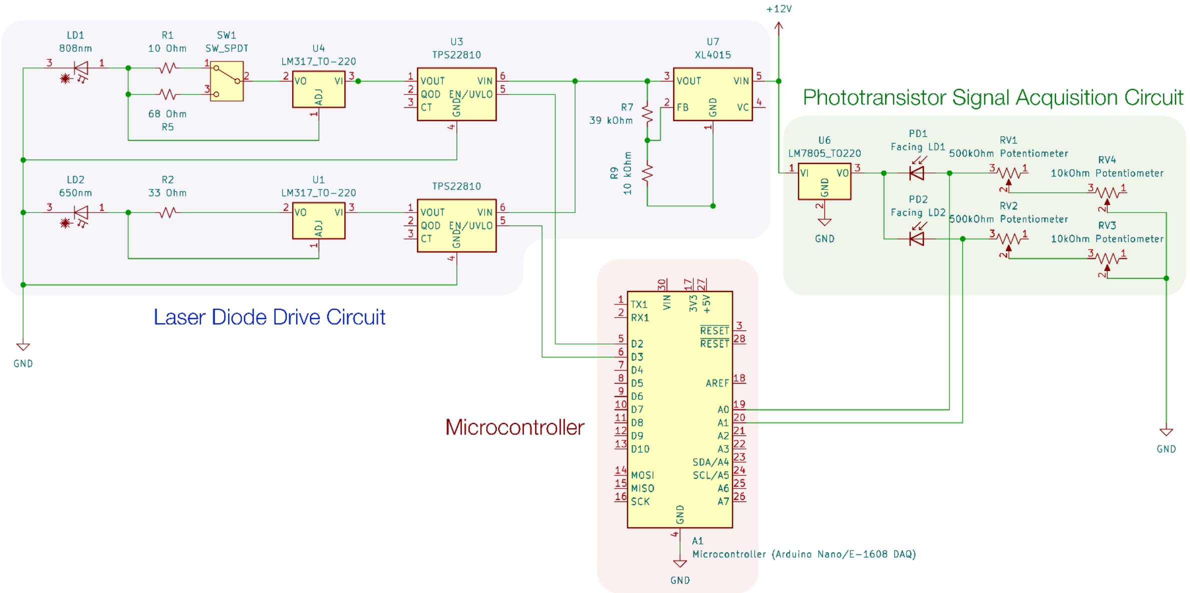


***Figure S2:*** *(A) Electrical Schematics of* ***(A)*** *Sensor v1.0, and* ***(B)*** *Sensor v1.1.*
