## Supplementary material for "Development and Validation of a Continuous Real-Time Optical Sensor for Indocyanine Green Clearance Measurement During Ex-Vivo Perfusion of Human Livers": Figure S3

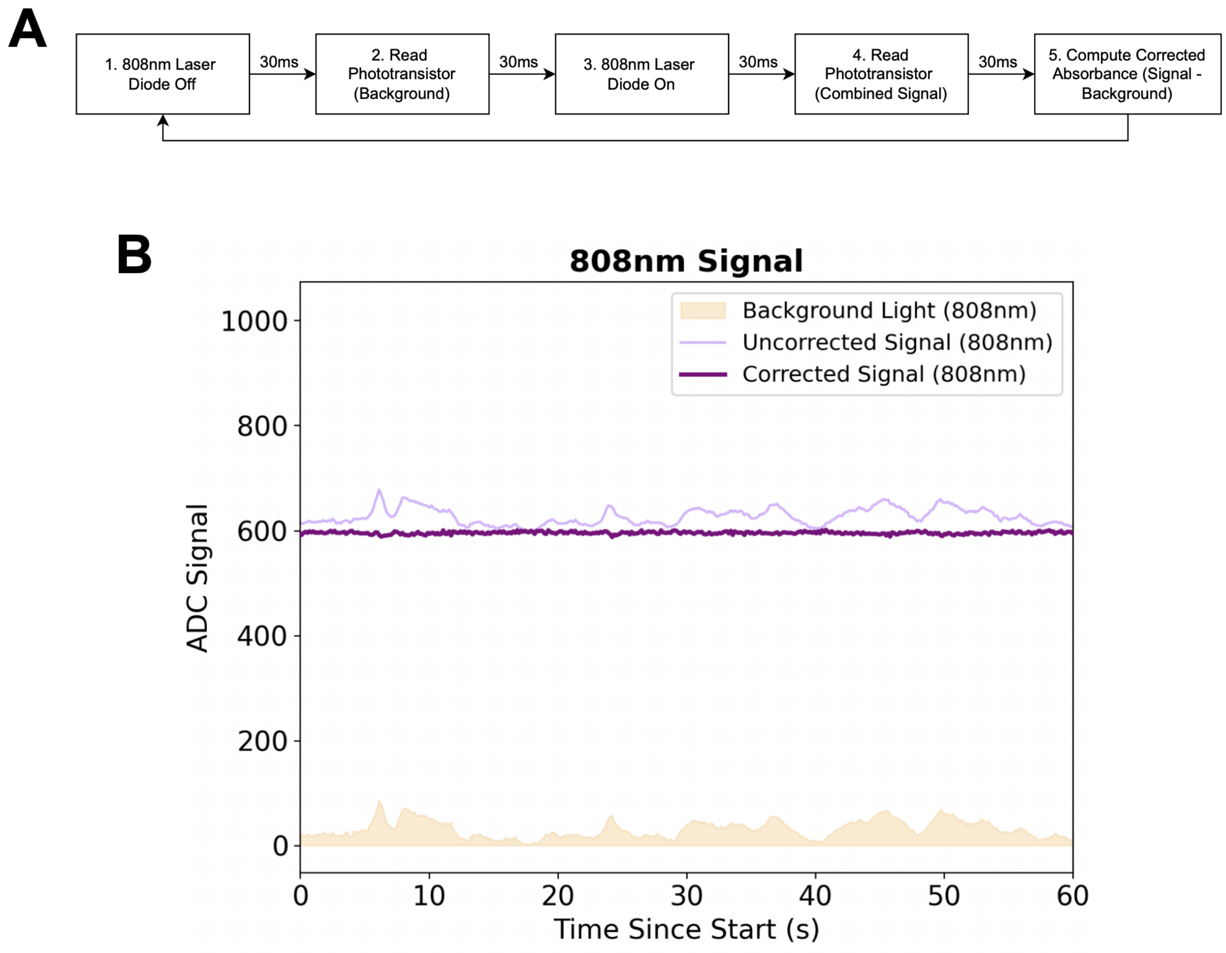


***Figure S3: Optical signal acquisition algorithm. (A)*** *Stepwise acquisition sequence implemented to isolate ICG-specific absorbance and minimise ambient noise. Each 120 ms measurement cycle consisted of five stages: (1) laser diode off for background preparation, (2) read background phototransistor signal, (3) laser diode on to illuminate the perfusate, (4) read combined signal, and (5) compute corrected absorbance (Δ808 = signal − background). This blinking and subtraction approach effectively removed ambient light interference and ensured stable, noise-free absorbance measurements.* ***(B)*** *808 nm signal showing raw, background, and corrected phototransistor readings.*
