## Supplementary material for "Development and Validation of a Continuous Real-Time Optical Sensor for Indocyanine Green Clearance Measurement During Ex-Vivo Perfusion of Human Livers": Figure S4

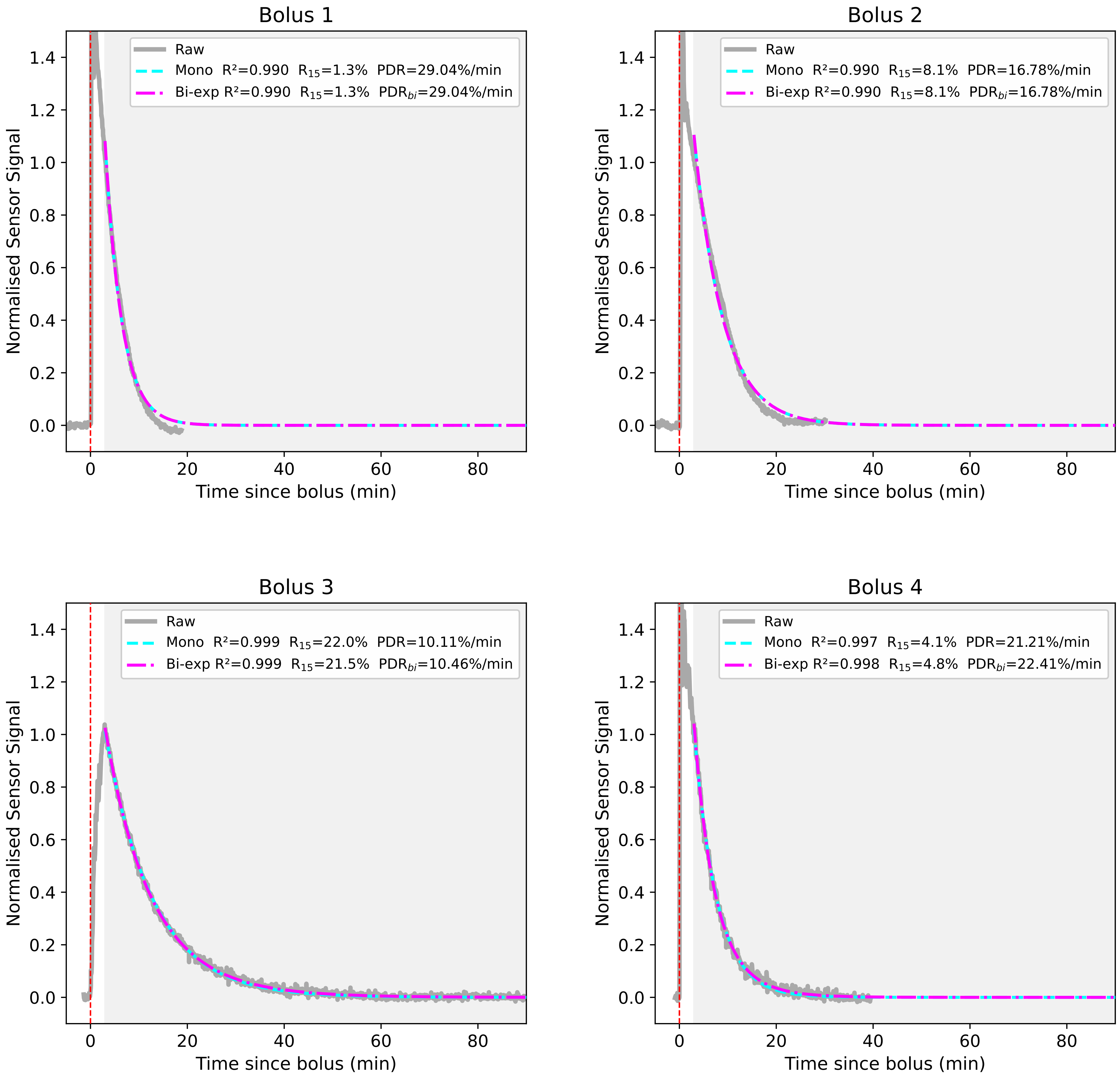
Bolus 1
Raw
Mono $R^2$=0.990 $R_{15}$=1.3% PDR=29.04%/min
Bi-exp $R^2$=0.990 $R_{15}$=1.3% $PDR_{bi}$=29.04%/min
Normalised Sensor Signal
Time since bolus (min)
Bolus 2
Raw
Mono $R^2$=0.990 $R_{15}$=8.1% PDR=16.78%/min
Bi-exp $R^2$=0.990 $R_{15}$=8.1% $PDR_{bi}$=16.78%/min
Normalised Sensor Signal
Time since bolus (min)
Bolus 3
Raw
Mono $R^2$=0.999 $R_{15}$=22.0% PDR=10.11%/min
Bi-exp $R^2$=0.999 $R_{15}$=21.5% $PDR_{bi}$=10.46%/min
Normalised Sensor Signal
Time since bolus (min)
Bolus 4
Raw
Mono $R^2$=0.997 $R_{15}$=4.1% PDR=21.21%/min
Bi-exp $R^2$=0.998 $R_{15}$=4.8% $PDR_{bi}$=22.41%/min
Normalised Sensor Signal
Time since bolus (min)

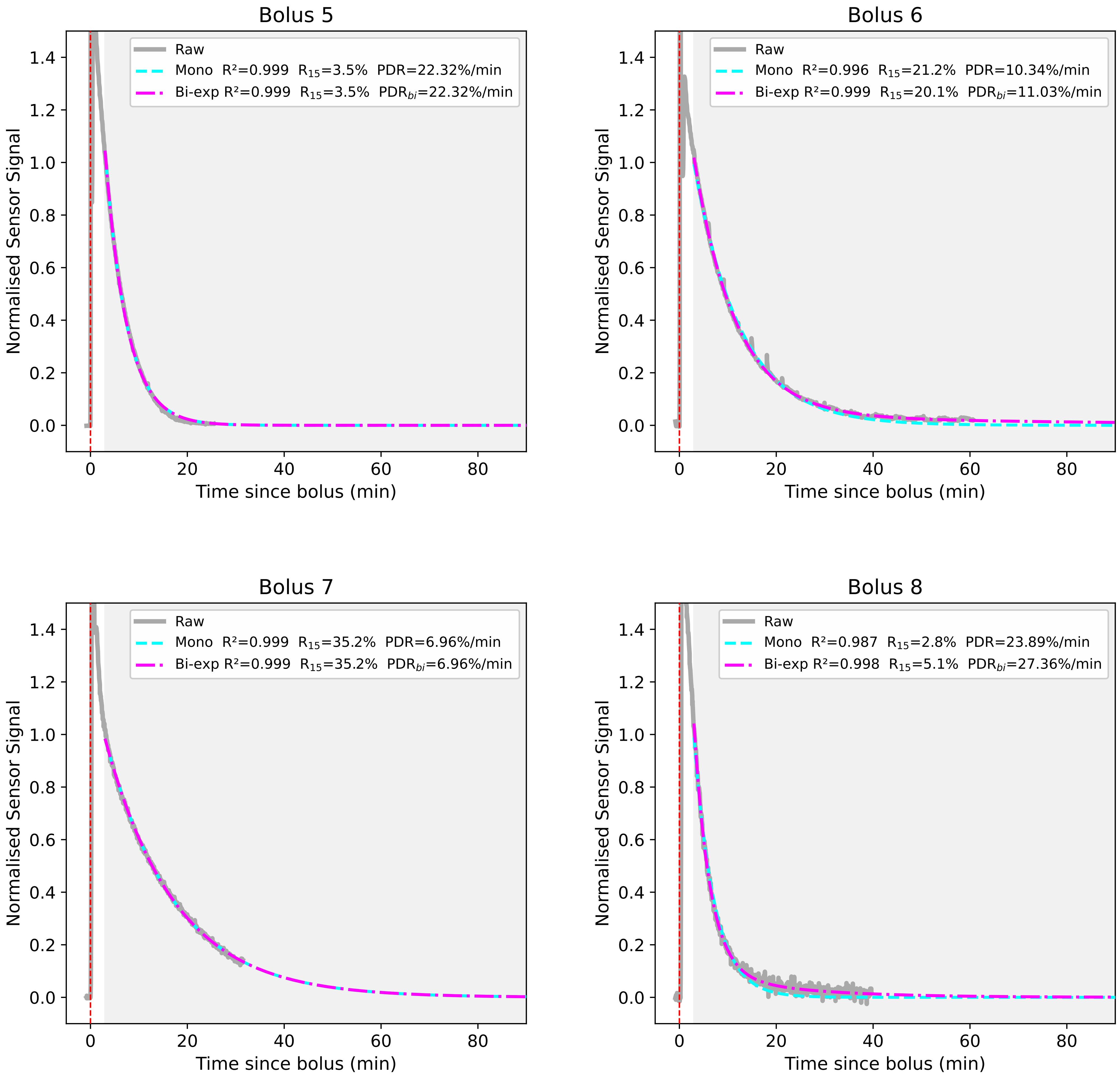
Bolus 5
Raw
Mono $R^2$=0.999 $R_{15}$=3.5% PDR=22.32%/min
Bi-exp $R^2$=0.999 $R_{15}$=3.5% $PDR_{bi}$=22.32%/min
Normalised Sensor Signal
Time since bolus (min)
Bolus 6
Raw
Mono $R^2$=0.996 $R_{15}$=21.2% PDR=10.34%/min
Bi-exp $R^2$=0.999 $R_{15}$=20.1% $PDR_{bi}$=11.03%/min
Normalised Sensor Signal
Time since bolus (min)
Bolus 7
Raw
Mono $R^2$=0.999 $R_{15}$=35.2% PDR=6.96%/min
Bi-exp $R^2$=0.999 $R_{15}$=35.2% $PDR_{bi}$=6.96%/min
Normalised Sensor Signal
Time since bolus (min)
Bolus 8
Raw
Mono $R^2$=0.987 $R_{15}$=2.8% PDR=23.89%/min
Bi-exp $R^2$=0.998 $R_{15}$=5.1% $PDR_{bi}$=27.36%/min
Normalised Sensor Signal
Time since bolus (min)

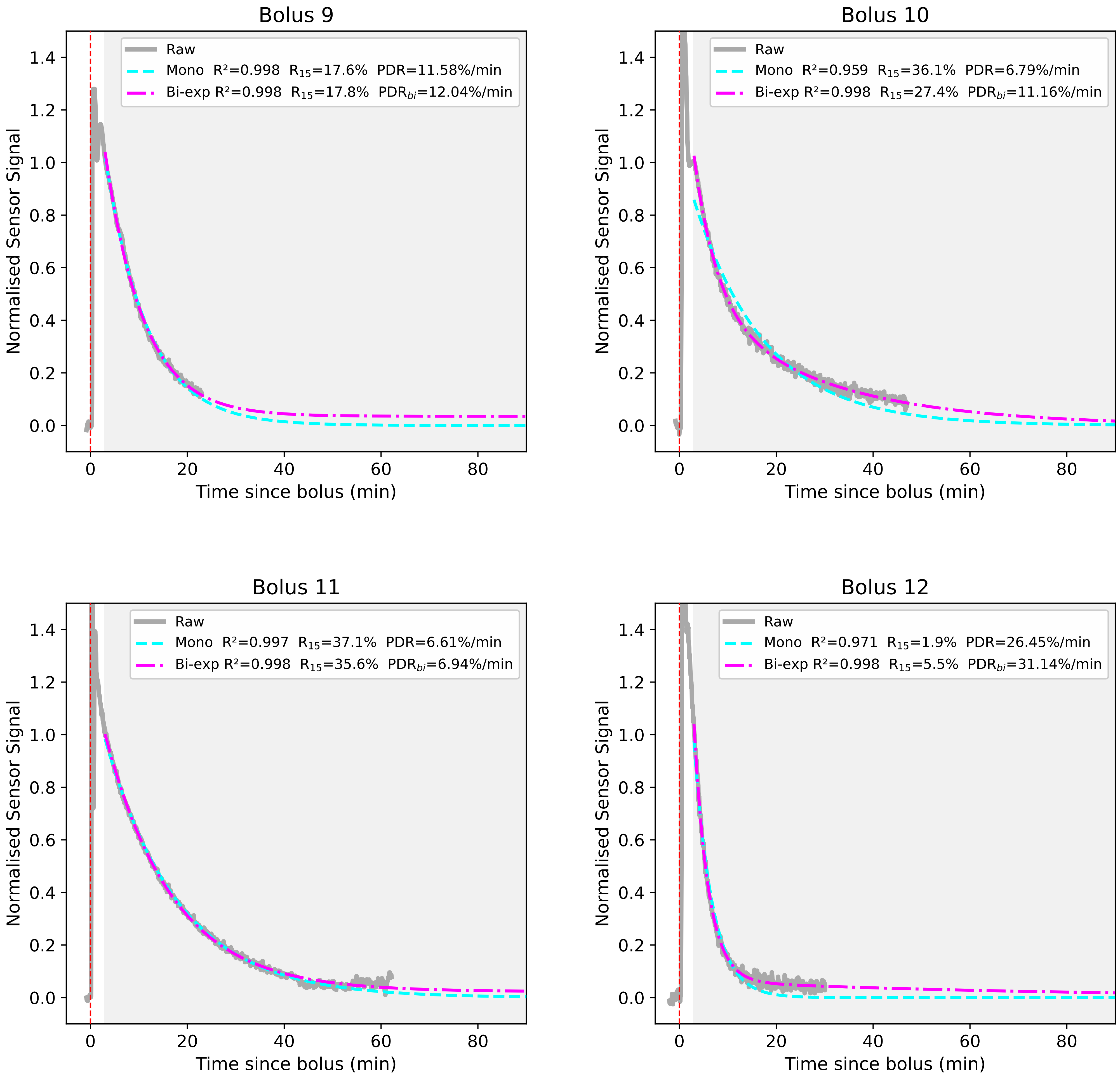
Bolus 9
Raw
Mono $R^2$=0.998 $R_{15}$=17.6% PDR=11.58%/min
Bi-exp $R^2$=0.998 $R_{15}$=17.8% $PDR_{bi}$=12.04%/min
Bolus 10
Raw
Mono $R^2$=0.959 $R_{15}$=36.1% PDR=6.79%/min
Bi-exp $R^2$=0.998 $R_{15}$=27.4% $PDR_{bi}$=11.16%/min
Bolus 11
Raw
Mono $R^2$=0.997 $R_{15}$=37.1% PDR=6.61%/min
Bi-exp $R^2$=0.998 $R_{15}$=35.6% $PDR_{bi}$=6.94%/min
Bolus 12
Raw
Mono $R^2$=0.971 $R_{15}$=1.9% PDR=26.45%/min
Bi-exp $R^2$=0.998 $R_{15}$=5.5% $PDR_{bi}$=31.14%/min
Normalised Sensor Signal
Time since bolus (min)
0.0
0.2
0.4
0.6
0.8
1.0
1.2
1.4
0
20
40
60
80

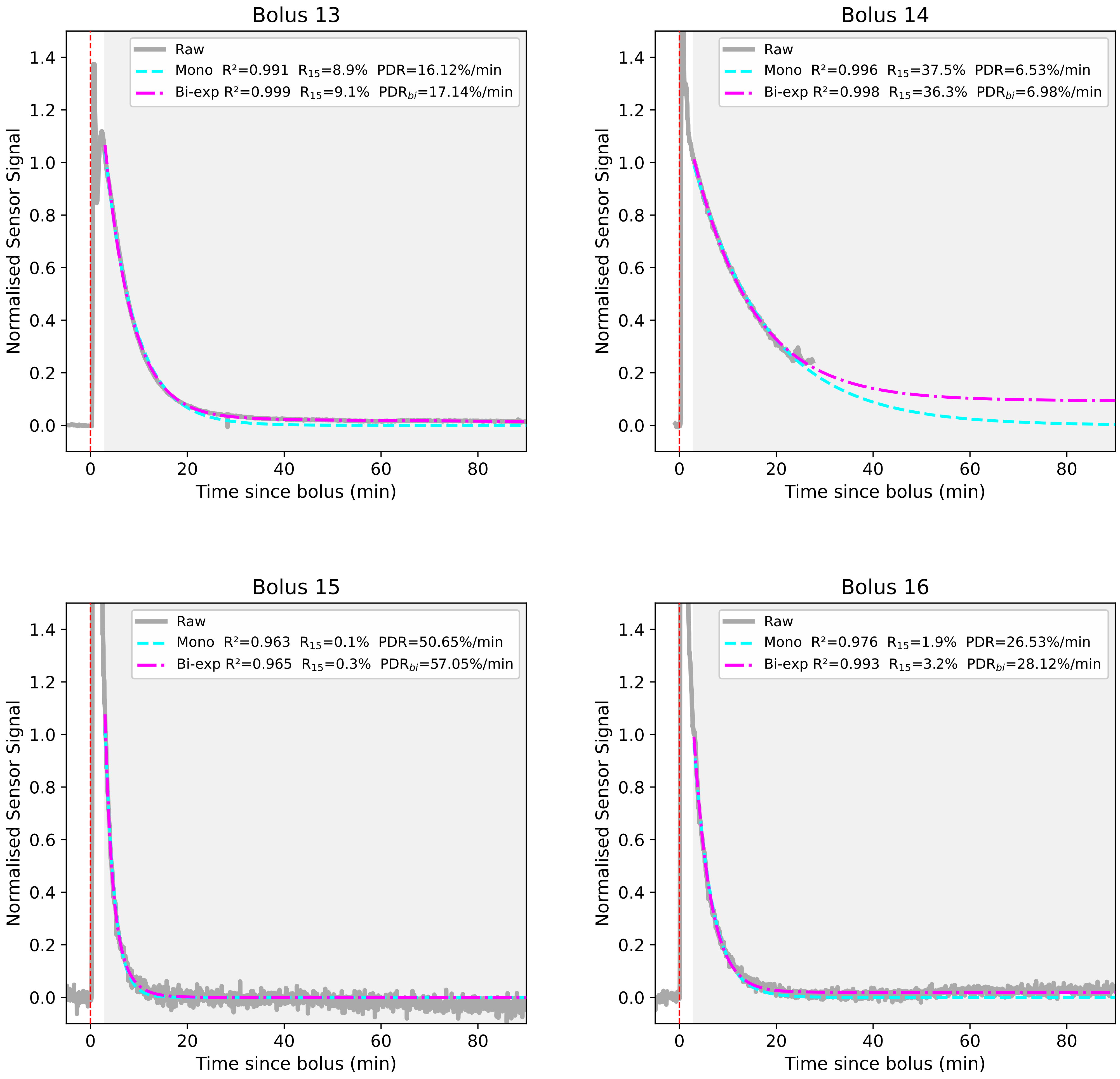

Bolus 13
Raw
Mono R²=0.991 R15=8.9% PDR=16.12%/min
Bi-exp R²=0.999 R15=9.1% PDRbi=17.14%/min
Normalised Sensor Signal
Time since bolus (min)
Bolus 14
Raw
Mono R²=0.996 R15=37.5% PDR=6.53%/min
Bi-exp R²=0.998 R15=36.3% PDRbi=6.98%/min
Normalised Sensor Signal
Time since bolus (min)
Bolus 15
Raw
Mono R²=0.963 R15=0.1% PDR=50.65%/min
Bi-exp R²=0.965 R15=0.3% PDRbi=57.05%/min
Normalised Sensor Signal
Time since bolus (min)
Bolus 16
Raw
Mono R²=0.976 R15=1.9% PDR=26.53%/min
Bi-exp R²=0.993 R15=3.2% PDRbi=28.12%/min
Normalised Sensor Signal
Time since bolus (min)

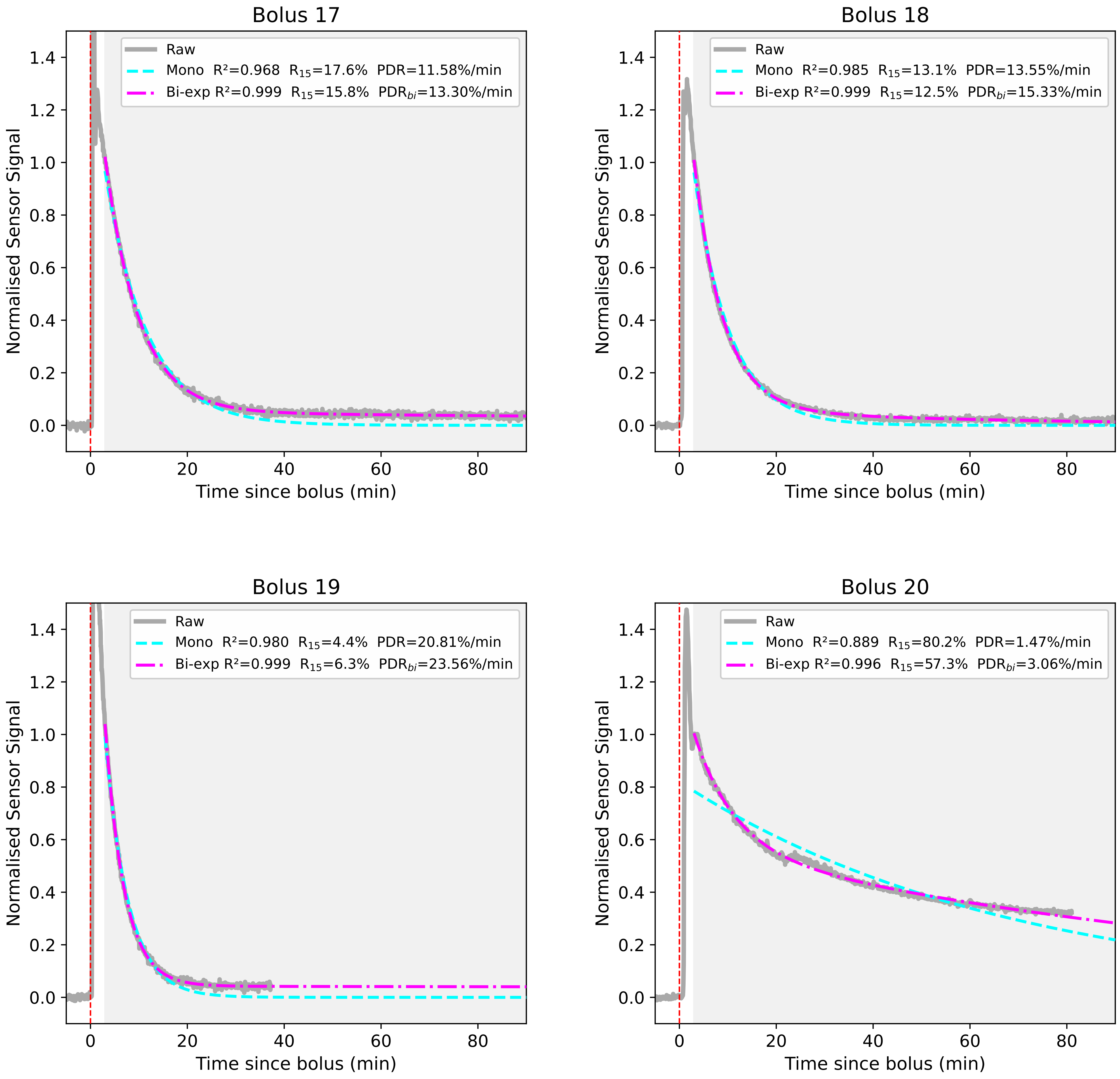
Bolus 17
Raw
Mono R²=0.968 R15=17.6% PDR=11.58%/min
Bi-exp R²=0.999 R15=15.8% PDRbi=13.30%/min
Bolus 18
Raw
Mono R²=0.985 R15=13.1% PDR=13.55%/min
Bi-exp R²=0.999 R15=12.5% PDRbi=15.33%/min
Bolus 19
Raw
Mono R²=0.980 R15=4.4% PDR=20.81%/min
Bi-exp R²=0.999 R15=6.3% PDRbi=23.56%/min
Bolus 20
Raw
Mono R²=0.889 R15=80.2% PDR=1.47%/min
Bi-exp R²=0.996 R15=57.3% PDRbi=3.06%/min
Normalised Sensor Signal
Time since bolus (min)

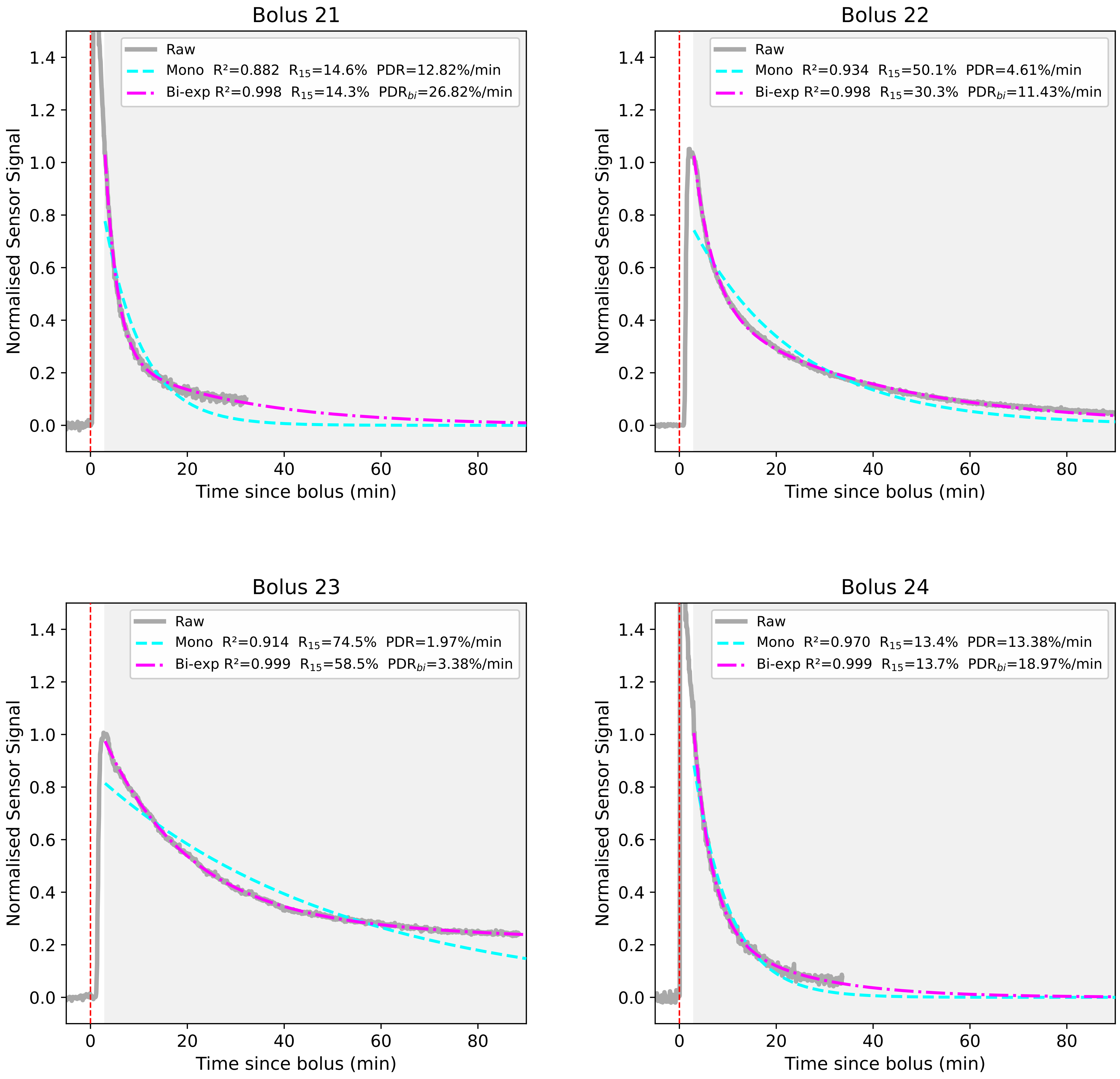

Bolus 21
Raw
Mono R²=0.882 R15=14.6% PDR=12.82%/min
Bi-exp R²=0.998 R15=14.3% PDRbi=26.82%/min
Normalised Sensor Signal
Time since bolus (min)
Bolus 22
Raw
Mono R²=0.934 R15=50.1% PDR=4.61%/min
Bi-exp R²=0.998 R15=30.3% PDRbi=11.43%/min
Normalised Sensor Signal
Time since bolus (min)
Bolus 23
Raw
Mono R²=0.914 R15=74.5% PDR=1.97%/min
Bi-exp R²=0.999 R15=58.5% PDRbi=3.38%/min
Normalised Sensor Signal
Time since bolus (min)
Bolus 24
Raw
Mono R²=0.970 R15=13.4% PDR=13.38%/min
Bi-exp R²=0.999 R15=13.7% PDRbi=18.97%/min
Normalised Sensor Signal
Time since bolus (min)

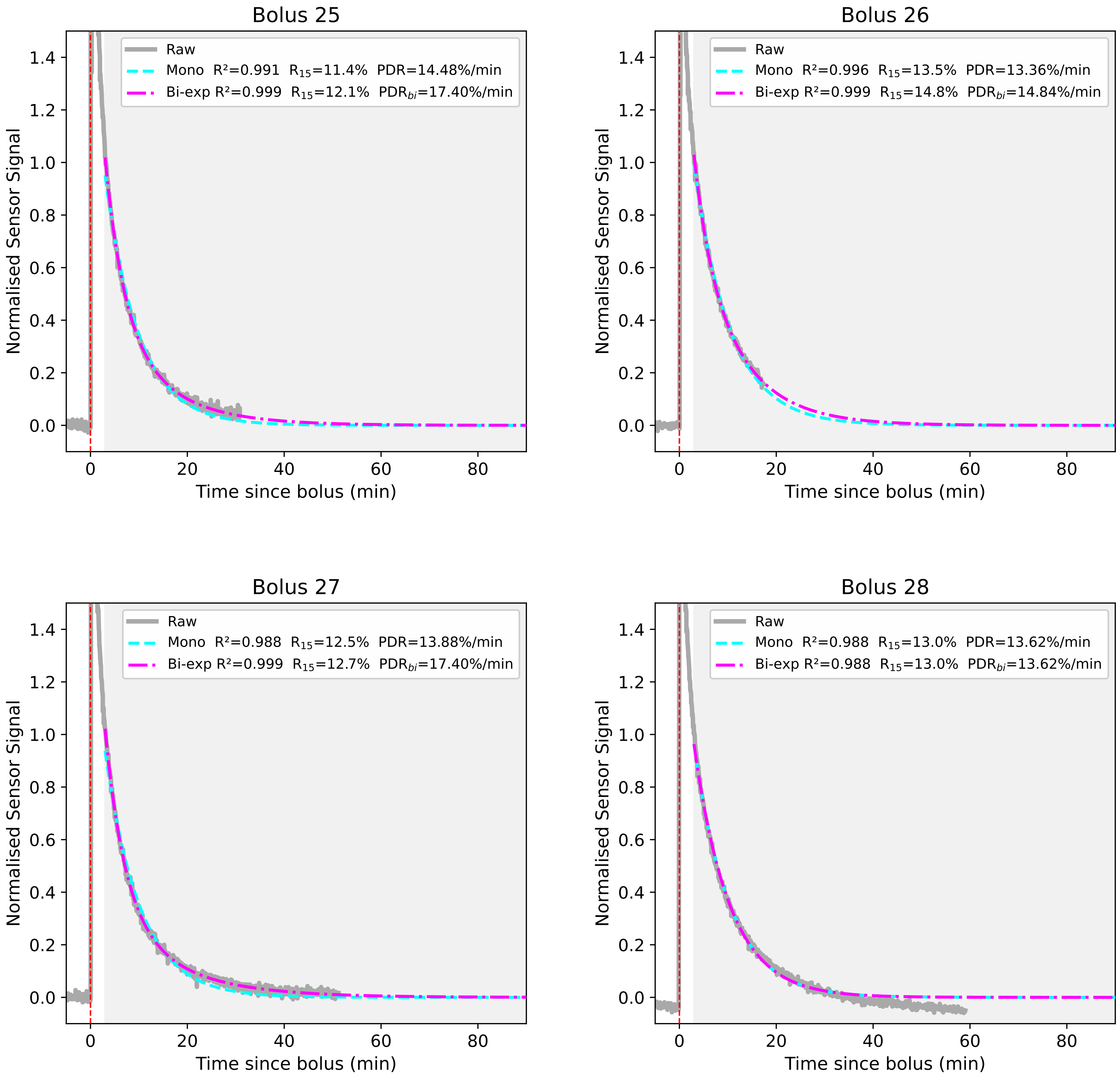

Bolus 25
Raw
Mono R²=0.991 R15=11.4% PDR=14.48%/min
Bi-exp R²=0.999 R15=12.1% PDRbi=17.40%/min
Normalised Sensor Signal
Time since bolus (min)
Bolus 26
Raw
Mono R²=0.996 R15=13.5% PDR=13.36%/min
Bi-exp R²=0.999 R15=14.8% PDRbi=14.84%/min
Normalised Sensor Signal
Time since bolus (min)
Bolus 27
Raw
Mono R²=0.988 R15=12.5% PDR=13.88%/min
Bi-exp R²=0.999 R15=12.7% PDRbi=17.40%/min
Normalised Sensor Signal
Time since bolus (min)
Bolus 28
Raw
Mono R²=0.988 R15=13.0% PDR=13.62%/min
Bi-exp R²=0.988 R15=13.0% PDRbi=13.62%/min
Normalised Sensor Signal
Time since bolus (min)

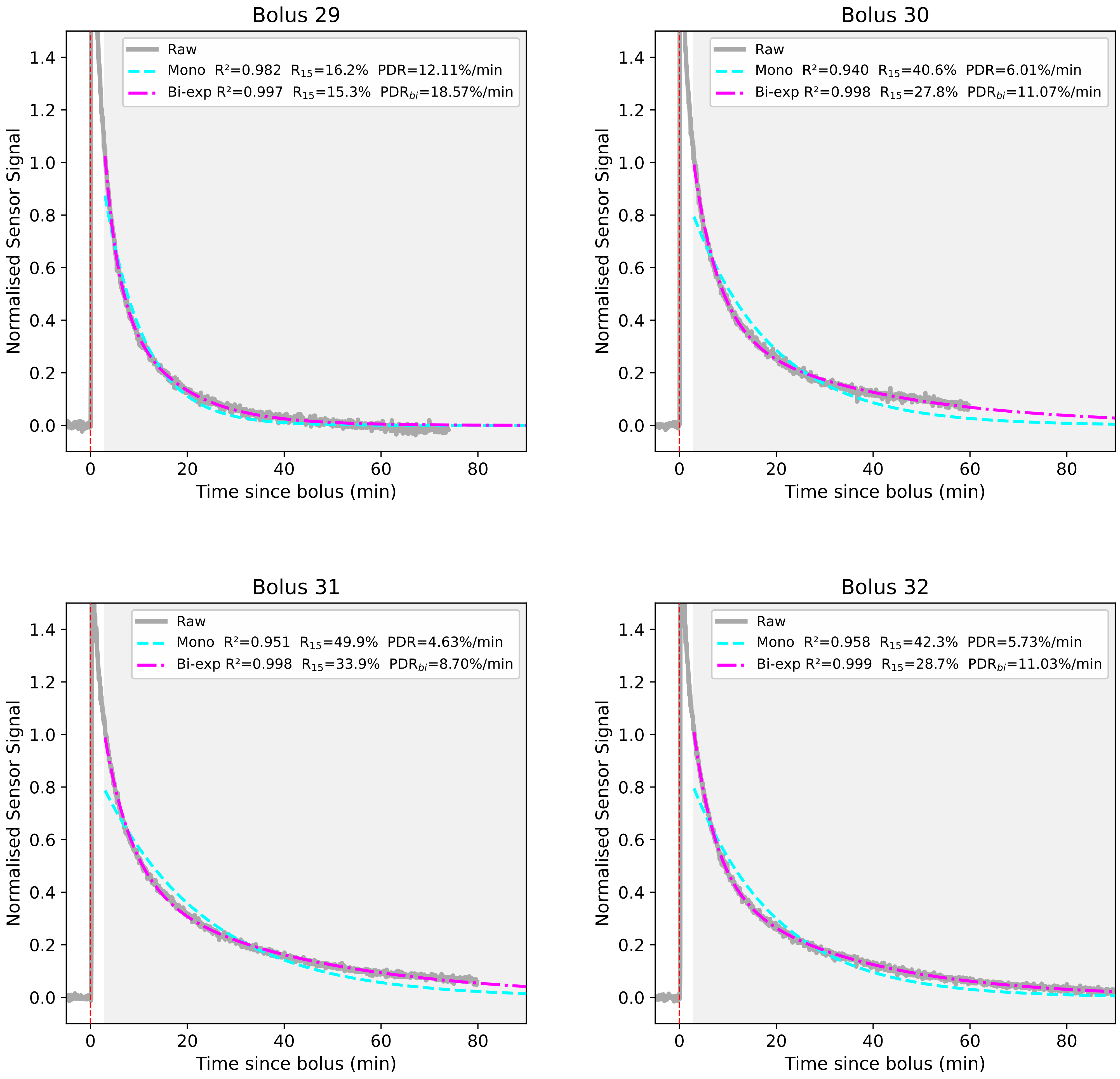
Bolus 29
Raw
Mono R²=0.982 R15=16.2% PDR=12.11%/min
Bi-exp R²=0.997 R15=15.3% PDRbi=18.57%/min
Normalised Sensor Signal
Time since bolus (min)
Bolus 30
Raw
Mono R²=0.940 R15=40.6% PDR=6.01%/min
Bi-exp R²=0.998 R15=27.8% PDRbi=11.07%/min
Normalised Sensor Signal
Time since bolus (min)
Bolus 31
Raw
Mono R²=0.951 R15=49.9% PDR=4.63%/min
Bi-exp R²=0.998 R15=33.9% PDRbi=8.70%/min
Normalised Sensor Signal
Time since bolus (min)
Bolus 32
Raw
Mono R²=0.958 R15=42.3% PDR=5.73%/min
Bi-exp R²=0.999 R15=28.7% PDRbi=11.03%/min
Normalised Sensor Signal
Time since bolus (min)

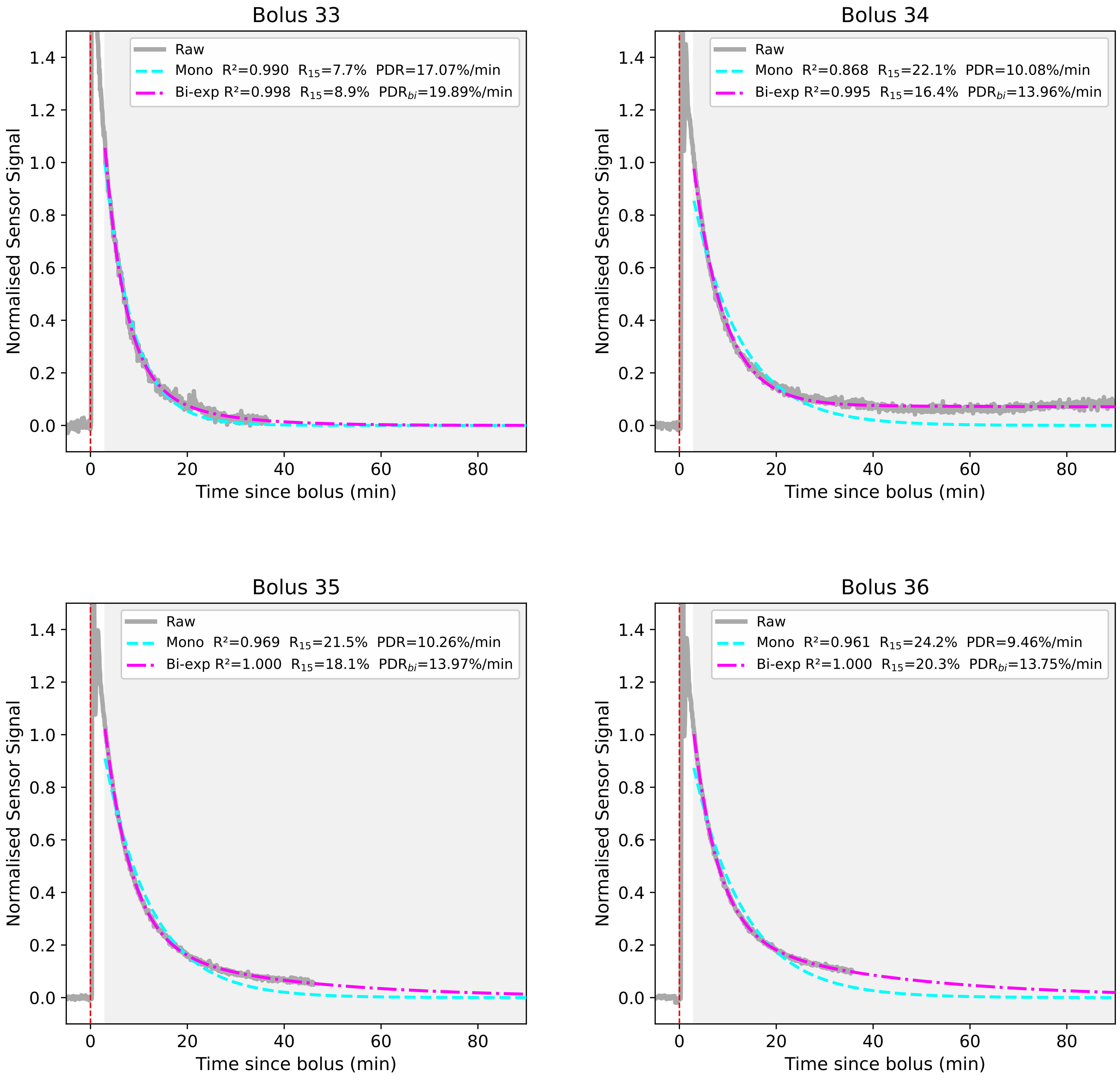
Bolus 33
Raw
Mono R²=0.990 R15=7.7% PDR=17.07%/min
Bi-exp R²=0.998 R15=8.9% PDRbi=19.89%/min
Normalised Sensor Signal
Time since bolus (min)
Bolus 34
Raw
Mono R²=0.868 R15=22.1% PDR=10.08%/min
Bi-exp R²=0.995 R15=16.4% PDRbi=13.96%/min
Normalised Sensor Signal
Time since bolus (min)
Bolus 35
Raw
Mono R²=0.969 R15=21.5% PDR=10.26%/min
Bi-exp R²=1.000 R15=18.1% PDRbi=13.97%/min
Normalised Sensor Signal
Time since bolus (min)
Bolus 36
Raw
Mono R²=0.961 R15=24.2% PDR=9.46%/min
Bi-exp R²=1.000 R15=20.3% PDRbi=13.75%/min
Normalised Sensor Signal
Time since bolus (min)

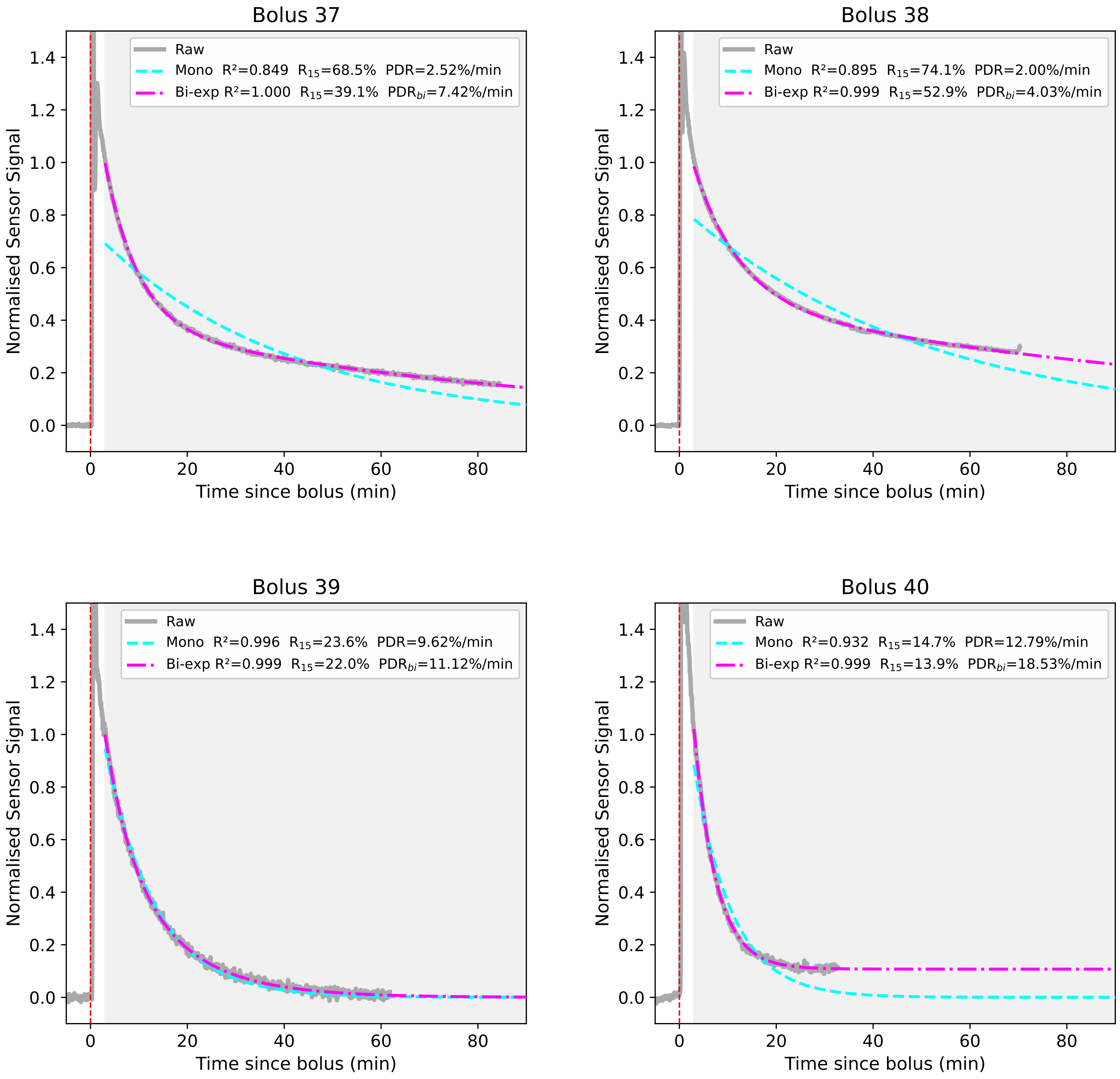

Bolus 37
Raw
Mono R²=0.849 R15=68.5% PDR=2.52%/min
Bi-exp R²=1.000 R15=39.1% PDRbi=7.42%/min
Bolus 38
Raw
Mono R²=0.895 R15=74.1% PDR=2.00%/min
Bi-exp R²=0.999 R15=52.9% PDRbi=4.03%/min
Bolus 39
Raw
Mono R²=0.996 R15=23.6% PDR=9.62%/min
Bi-exp R²=0.999 R15=22.0% PDRbi=11.12%/min
Bolus 40
Raw
Mono R²=0.932 R15=14.7% PDR=12.79%/min
Bi-exp R²=0.999 R15=13.9% PDRbi=18.53%/min
Normalised Sensor Signal
Time since bolus (min)

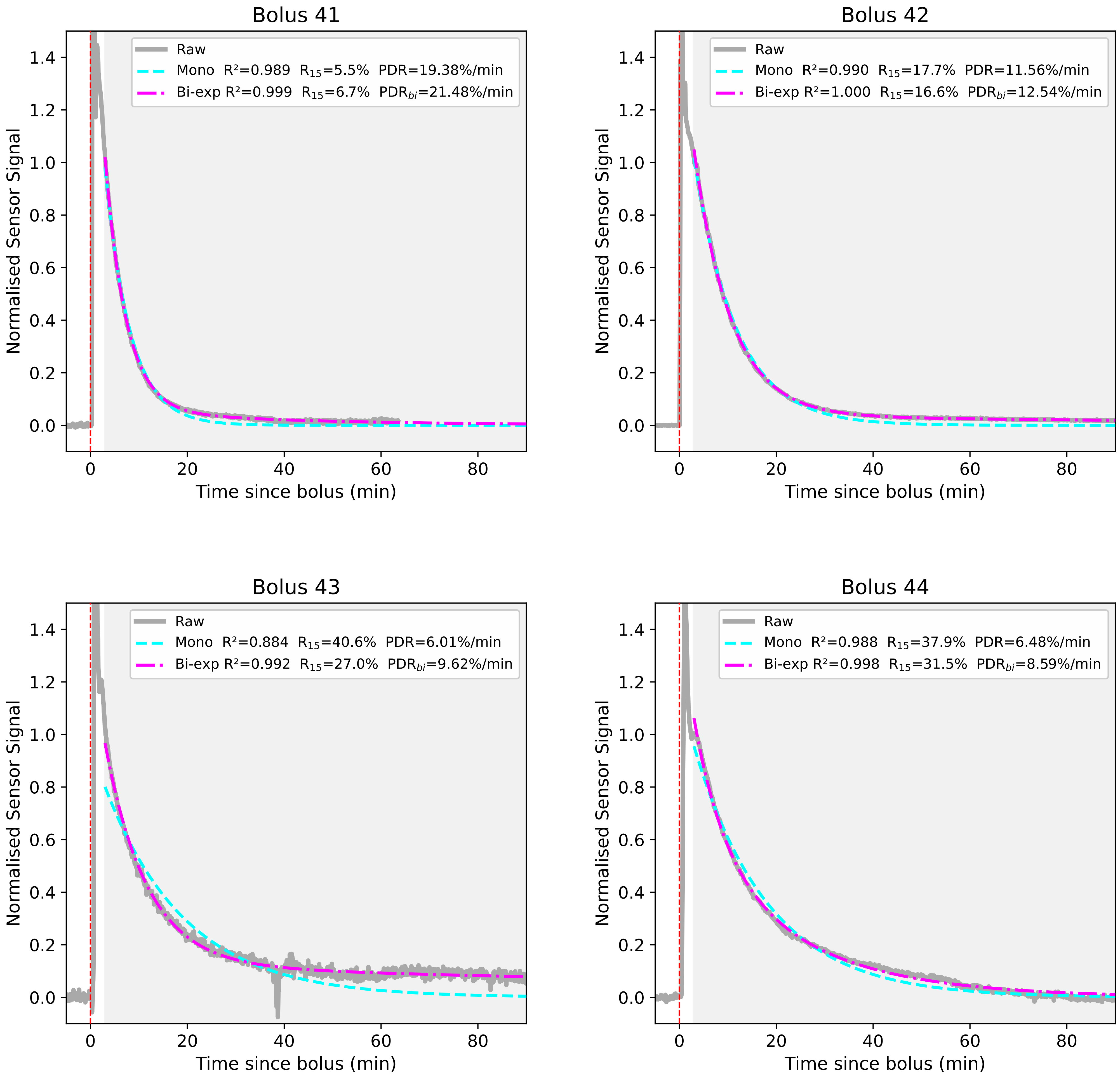

Bolus 41
Raw
Mono R²=0.989 R15=5.5% PDR=19.38%/min
Bi-exp R²=0.999 R15=6.7% PDRbi=21.48%/min
Normalised Sensor Signal
Time since bolus (min)
Bolus 42
Raw
Mono R²=0.990 R15=17.7% PDR=11.56%/min
Bi-exp R²=1.000 R15=16.6% PDRbi=12.54%/min
Normalised Sensor Signal
Time since bolus (min)
Bolus 43
Raw
Mono R²=0.884 R15=40.6% PDR=6.01%/min
Bi-exp R²=0.992 R15=27.0% PDRbi=9.62%/min
Normalised Sensor Signal
Time since bolus (min)
Bolus 44
Raw
Mono R²=0.988 R15=37.9% PDR=6.48%/min
Bi-exp R²=0.998 R15=31.5% PDRbi=8.59%/min
Normalised Sensor Signal
Time since bolus (min)

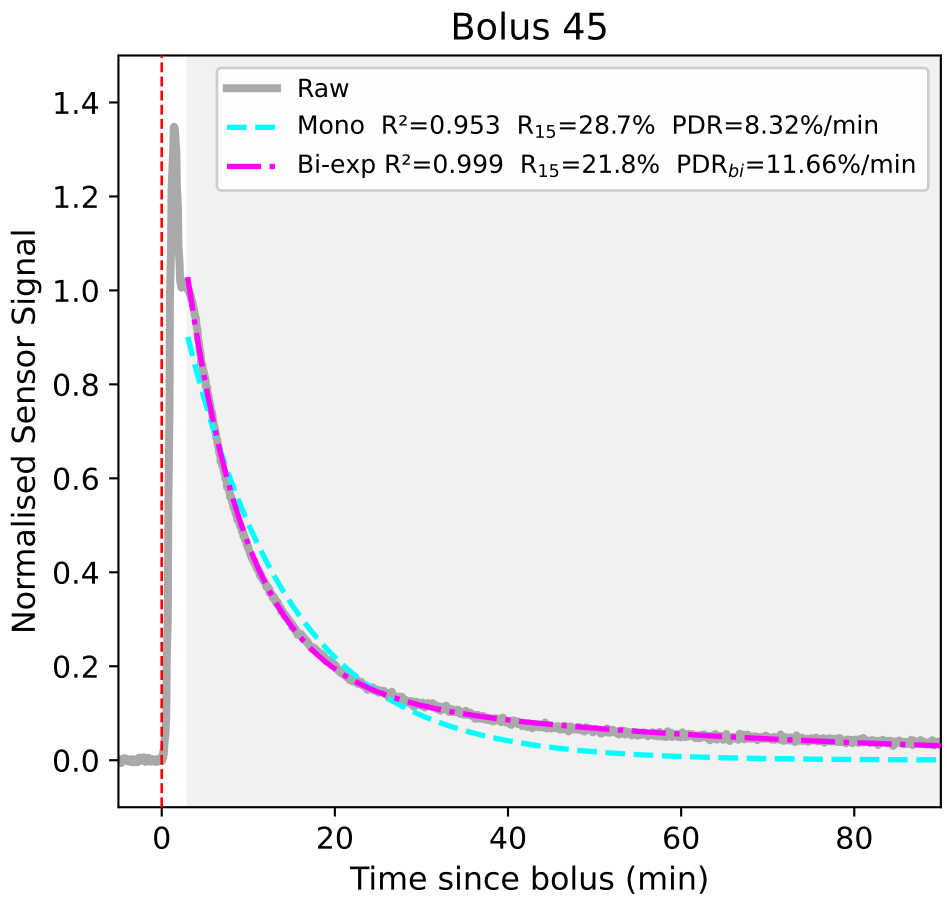


*Figure S4: Compiled 45 indocyanine green (ICG) clearance boluses with mono- and bi-exponential fitted models (baseline C fixed to pre-bolus baseline). Each plot shows normalised sensor data with corresponding R2, R15 and PDR/PDRbi values. Across boluses, R2 summaries were: bi-exponential - mean 0.997, median 0.999, range 0.965–0.9997; mono-exponential - mean 0.964, median 0.982, range 0.849–0.999.*
